# Multi-enhancer regulatory hubs control Th17 cell identity

**DOI:** 10.64898/2026.04.02.715458

**Authors:** Keith Siklenka, Chuangchuang Zhang, Liqing Li, Morgan Parker, Naren Mehta, Alejandro Barrera, Revathy Venukuttan, Gregory E Crawford, Charles A Gersbach, Maria Ciofani, Timothy E Reddy

## Abstract

Between one and two million noncoding regulatory elements have been described across mammalian genomes, but determining their functional role remains a challenge. To address this gap, we measure the regulatory potential of open chromatin regions (OCRs) in five closely related mouse CD4+ T cell subsets with ATAC-STARR-seq, then map endogenous enhancer function and three-dimensional contacts in Th17 cells with pooled CRISPR-based noncoding screens and high-resolution Micro-C. Of all CD4+ T cell OCRs, approximately 25% demonstrate largely subset-shared regulatory activity, though we identify subset-restricted active elements distinguishable by their sequence features. In Th17 cells, we reveal a set of core regulatory OCRs essential for subset polarization at the *Batf*, *Rorc(t)* and *Il17a/f* loci, and confirm their requirement for cell identity *in vivo*. At these three loci, we resolve nested yet selective enhancer-to-enhancer and enhancer-to-promoter physical interactions that converge into multi-enhancer hubs. Importantly, these hubs contain many of the strongest functional elements. Perturbation of a single enhancer within the *Batf* locus selectively disrupts hub contacts and *Batf* expression in Th17 cells, recapitulating the genome-wide transcriptional and chromatin accessibility signatures of a germline knockout. Together, this work assigns regulatory activity to accessible chromatin across CD4+ T cell subsets and couples the function of a set of essential elements to selective three-dimensional contacts in Th17 cells.

## Introduction

Genome-wide maps have nominated hundreds of thousands of candidate regulatory elements across mammalian cell types based on their associated chromatin features^1–3^. Yet there remains a critical disconnect between regulatory annotation and regulatory function. High-throughput functional assays such as massively parallel reporter assays (MPRAs) and self-transcribing active regulatory region sequencing (STARR-seq) can evaluate sequence regulatory potential in an episomal context^4,5^, while pooled CRISPR screens can measure endogenous function in the native chromatin context^6^, and chromatin conformation capture resolves the 3D architecture in which elements act^7,8^. Most studies evaluate these properties in isolation, often resulting in discordant findings that are difficult to interpret.

MPRA and STARR-seq have successfully mapped regulatory activity in immortalized cell lines^4,9–12^ and select primary cells^13–15^, but how sequence-dependent regulatory potential operates across closely related cellular contexts is unclear. Similarly, pooled CRISPR-based screens have largely focused on identifying key *trans*-acting regulators of primary T cell function^16–18^, leaving the underlying network of *cis*-regulatory elements largely uncharacterized^19–21^. Murine CD4+ T cell differentiation provides an ideal system to bridge these gaps. Upon activation and specific cytokine signalling, naive CD4 T cells (Tn) undergo major transcriptional and structural remodeling as they polarize into functionally distinct subsets, including T helper 1 (Th1), Th2, Th17 and regulatory T cells (Treg). This process requires genome-scale coordination of cell-type-specific expression programs, yet the *cis*-regulatory networks that orchestrate these changes remain incompletely resolved.

Here, we show regulatory activity emerges from three layers of functional information: sequence-encoded regulatory potential, native chromatin context, and the three-dimensional architecture. To do so, we integrate large-scale episomal reporter assays, endogenous epigenetic perturbations, and high-resolution 3D genomics to map the regulatory architecture of T cell identity. By adapting ATAC-STARR-seq across five distinct mouse T cell subsets, we systematically measure all open chromatin DNA for intrinsic sequence regulatory potential. We show that regulatory activity across OCRs is largely shared between subsets and is independent of the endogenous chromatin state. We determine the degree of which regulatory activity is biased towards an individual subset, and how that activity is characterized by cell-type specific transcription factor motifs. Through pooled CRISPR screens in Th17 cells we perturb thousands of open chromatin regions to identify a set of critical *cis*-regulatory elements at the *Batf*, *Rorc* and *Il17* loci that are functionally necessary for Th17 cell identity. By integrating high-resolution region capture micro-C (RCMC) we associate functional enhancers with hub-like, multi-element physical interactions that coordinate regulation of their gene targets. Strikingly, we demonstrate how a single epigenetic perturbation of a distal *Batf* enhancer disrupts 3D chromatin contacts leading to downstream genome-wide regulatory impacts comparable to those of a complete knockout, severely impairing Th17 cell phenotypes. Together, this work establishes how sequence encoded regulatory potential, enhancer function, and 3D chromatin topology collaborate to direct T cell identity.

## Results

### Defining candidate regulatory elements in mouse CD4+ T cells

To define the gene regulatory landscape of mouse CD4+ T cells (Fig. 1a), we first mapped genome-wide chromatin accessibility with ATAC-seq in Th0 (activated naive T cells), Th1, Th2, Th17, and Treg cells, identifying 132,082 OCRs across all subsets (Supplemental Table 1). Through comparative analysis, we show extensive subset-specific chromatin remodelling occurs during T cell polarization consistent with large scale *cis*-regulatory atlases of the mouse immune system^2^. We identified 43 to 66 thousand (31-48%) differentially accessible OCRs in each subset relative to the Th0 state (Supplementary Table 1), and 2 to 14 thousand (2-10%) unique OCRs per subset (Fig. 1c, Supplementary Fig 1b). To measure regulatory activity of these CD4 OCRs across each T cell subset, we generated a unified ATAC-STARR-seq DNA library (input) by pooling accessible fragments recovered from FACS purified cells (see Methods). Deep sequencing of this library confirmed near complete capture of all identified CD4+ OCRs, achieving >100x unique-fragment coverage for 98% of peak regions (Supplementary Fig. 1a-c). The OCRs identified in this pooled input library served as the union-set of candidate regulatory elements for all downstream functional analysis (Supplementary Table 1).

**Figure 1.**
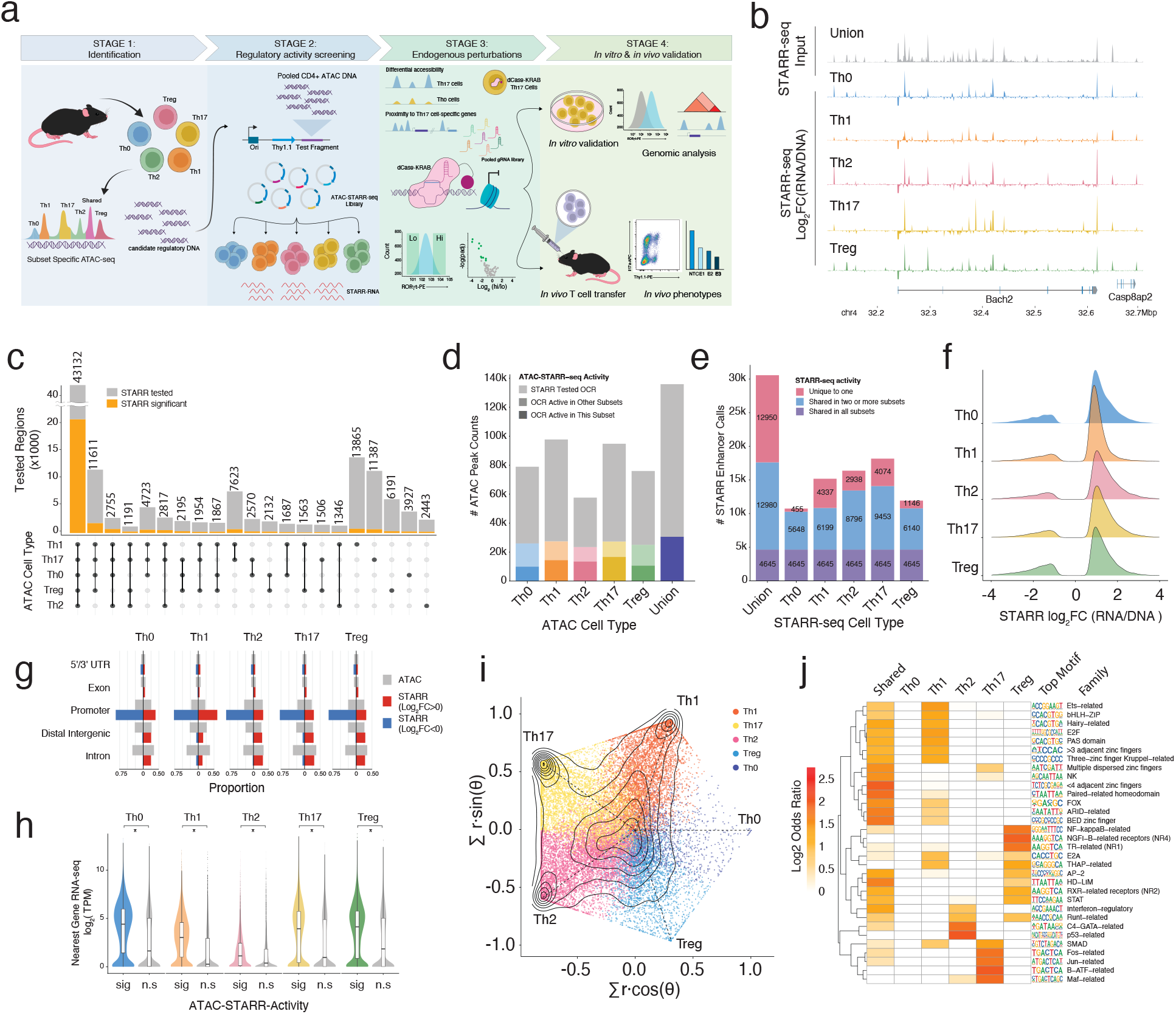
High-throughput reporter assays reveal shared and context-dependent enhancer activity in CD4 T cells. **a)** Schematic of project design. Stage 1: Identify candidate regulatory elements from in vitro derived Th0, Th1, Th2, Th17 and Treg CD4+ subsets with ATAC- seq. Stage 2: Pool all ATAC-seq fragments and test their regulatory potential across all CD4+ T cell subsets with ATAC-STARR-seq. Stage 3: Perform pooled CRISPR-based epigenome editing screens subset differentiation in Th17 cells by targeting OCRs with the most differential accessibility nearby Th17-relevant gene loci. Stage 4: Perform in vitro and in vivo validation of single enhancer perturbations in Th17 cells followed by flow cytometry readouts for cell phenotypes, and genome wide analyses of gene expression, chromatin accessibility and three- dimensional chromatin contacts. **b)** Genome browser track depicting the union ATAC-STARR-seq input library (grey) and STARR-seq activity score (Log2 CPM RNA/DNA; coloured) at the Bach2 locus for each subset. **c)** Upset plot of OCR intersections between all CD4 T cell subsets, with the proportion of STARR-seq active OCRs for each intersection highlighted in yellow. **d)** Stacked bar plots indicating the total number of OCRs tested in ATAC-STARR-seq and the proportion of OCRs with significant STARR-seq activity in that subset (dark colour), any subset (light colour) or tested but not significant (grey; fdr < 0.05). **e)** Total STARR-seq enhancer calls by subset (FDR < 0.05) with the proportion of active OCRs labelled by chromatin category (Shared in all subsets = purple; shared in two or more subsets = blue; unique to one subset = red) **f)** Effect sizes of significant enhancer calls for each cell type (Log2 RNA/DNA) **g)** Genome annotation of OCRs mapped by ATAC-seq (grey) or functional OCRs determined by STARR-seq (active = red; repressive = blue) for each subset. **h)** Violin plot showing distribution and median transcript abundance of genes nearest to OCRs categorized by ATAC-STARR-seq enhancer call significance for that subset (sig = STARR FDR < 0.05; n.s = non-significant; all p ∼ 0 for sig vs n.s, Mann-Whitney U Test) **i)** Radial plot of STARR-seq regulatory activity preference across CD4+ T cell subsets. Each axis represents a tested subset (Th0, Th1, Th2, Th17, Treg), and each point represents an active OCR with significant STARR-seq activity in at least one subset (FDR < 0.01). A weighted activity score (|Log2FC| x -log10(FDR)) of each OCR was calculated for each subset, normalized to sum to one across all subsets, and projected into 2D using angular coordinates. OCRs with STARR- seq activity biased toward a single subset fall near the corresponding spoke, while OCRs with equivalent activity across subsets fall near the center. Points are coloured by the subset in which the OCR exhibits its highest activity score. **j)** Odds ratio of top enriched motif families at subset-preferred OCRs. For each enhancer-preference group (subset-preferred, r = 0.85; subset-shared, r=0.3), odds ratios were computed for the preferred OCR set against all others. Top motif families are shown.

### Sequence-encoded regulatory activity is largely shared across CD4+ T cell subsets

We then introduced the ATAC-STARR-seq library into each T cell subset to quantify the regulatory activity of OCRs across these closely related but functionally distinct cells. This episomal assay evaluates DNA sequences independent of chromatin context, therefore we interpret assay signals as the sequence-encoded regulatory potential. Overall, we identified 30,575 active OCRs across all T cell subsets (∼8,000-18,000 active elements per subset), representing 23% of tested regions (Fig 1c-e; Supplementary Table 1). Most active OCRs were positive regulators of STARR-seq signal and mapped equally across genomic contexts, while a minority of regions acted as negative regulators and mapped primarily to promoters (Fig. 1f,g). Across all subsets, genes nearest active OCRs were expressed significantly higher than those near inactive OCRs (Fig. 1h).

To investigate whether active or repressive OCRs harbour distinct sequence features, we trained gapped k-mer support vector machines (gkmSVM)^22,23^ to distinguish active and repressive OCRs from inactive OCRs in each subset. Activated elements were classified with moderate accuracy (AUC 0.73-0.80), driven primarily by enrichments of CRE/ATF motifs (all subsets), AP-1/TRE (Th17), AT-rich homeodomain-like motifs (Treg), and GC-rich motifs matching KLF/Sp family (Th1) (Supplementary Fig. 2c,e). Repressive elements, however, were characterized almost entirely by AT and GC dense sequences rather than identifiable repressive TF binding motifs (Supplementary Fig. 2d,f).

We then assessed enhancer activity across subsets as it relates to chromatin accessibility in the native context of the cell. Despite large scale chromatin specification during T cell polarization, STARR-seq activity at CD4+ OCRs were strongly correlated across all T cell subsets (average Pearson r = 0.72; Supplementary Fig. 1h). As expected, the majority of active STARR signal corresponded to OCRs with subset-shared accessibility (Supplementary Fig. 1e). Strikingly, however, we identified robust STARR-seq signal across all tested T cells at OCRs with subset- unique accessibility. This latent regulatory potential occurred with stronger average effect sizes than the shared OCR categories (Supplementary Fig. 1g), highlighting how cell-type specific enhancer sequences may encode potent activity that is functionally restricted in different contexts. Consistent with this, we found no global correlation between the absolute magnitude of chromatin accessibility and STARR-seq regulatory activity (Supplementary Fig. 3b,c), and only a weak positive relationship with subset-polarization induced changes (r = 0.1 to 0.38, Supplementary Fig. 3a). Together, these results indicate that chromatin accessibility restricts, rather than creates, the regulatory activity encoded at STARR-seq positive regions.

### Subset-specific activity emerges amid shared elements

Within the shared regulatory landscape of CD4 T cell subsets, we identified distinct categories of elements with subset-specific STARR-seq activity. To resolve these regions, we computed a weighted effect size for OCRs across subsets and mapped them to a radial plot reflecting preference and magnitude (Fig. 1g; Supplementary Fig. 4; See Methods). The distribution of regulatory elements resulted in both a pronounced central density, indicating broadly shared activity, and radial densities for each subset, indicating specific activity (Fig. 1g). Subset-specific regulatory activity was strongly biased toward effector subsets (Th1, Th2, and Th17), whereas Treg-preferred elements were markedly underrepresented, and Th0-specific elements strongly depleted. These patterns suggest that the regulatory potential of a distinct class of CD4 T cell OCRs is governed by a sequence-encoded regulatory logic sensitive to subset-specific contexts.

To identify the sequence determinants of this subset-specificity, we compared transcription factor (TF) motif enrichment in the most subset-specific regulatory OCRs at the radial poles versus the most shared regions at the center (Supplementary Fig. 4c; Table 2). Elements with shared regulatory potential were enriched for diverse motifs representing many different classes of transcription factors (Fig. 1h). In contrast, the subset-specific active OCRs were enriched for motifs of TF families with defined roles that relate to each subset (Fig. 1h). For instance, the Th17 cell-preferential elements were enriched for AP-1^3,24,25^, SMAD^26^, and Maf-related^27^ transcription factors, key components of type 17 effector programming. Th2-preferential elements were enriched for GATA-family motifs, consistent with GATA3’s master regulatory role^28–30^, as well as Runt^31^ and IRF motifs^32,33^. The Treg-preferential elements were enriched for cytokine responsive motifs (NFkB and STAT), while the Th1-preferential elements were strongly enriched for Ets and Fox family members. To assess if this subset-preferred STARR-seq activity reflects differential transcription factor abundance, we examined expression of the enriched TF families across subsets, revealing broadly shared abundance (Supplementary Fig. 4e). Consistent with this, expression levels of all identified TF and epigenetic regulators were highly correlated between subsets (r = 0.46 – 0.92; Supplementary Fig. 4d).

Taken together, functional characterization of the accessible chromatin landscape in CD4 T cells with STARR-seq revealed a broadly shared regulatory potential that is overlaid by a distinct category of subset-specific regulatory elements. Because of similarities in overall TF abundance in these cells, the subset restricted activity at these elements implicates sequence-level features as the primary driver of this specificity.

### CRISPR-based characterization of functional Th17 cell regulatory elements

Although episomal reporter assays identified thousands of OCRs with regulatory potential, they cannot determine which elements are functionally required in their native chromatin context. To address this, we sought to identify OCRs with essential function in Th17 cell subset specification using CRISPR interference (CRISPRi). We reasoned that OCRs exhibiting the greatest increase in both accessibility and expression of neighbouring genes would represent the most potent subset-essential enhancers and designed a pooled CRISPRi screening library to target 7,000 of the top-ranked Th17 cell-induced OCRs. The final library consisted of 68,870 single guide RNAs (sgRNAs) targeting OCRs with approximately 10 sgRNAs per element, and 5% allocated to non-targeting controls (Supplementary Table 3, See Methods). We transduced the library at low MOI (∼0.3) into Th0 cells from mice expressing both dCas9-KRAB and an *Il17a* locus eGFP reporter (*Il17a*-eGFP). Following culture in Th17 cell-polarizing conditions, cells were sorted based on reporter expression, and functional regulators were identified by comparing sgRNA abundance in *Il17a*-eGFP-high vs *Il17a*-eGFP-low populations (Fig. 2a).

**Figure 2.**
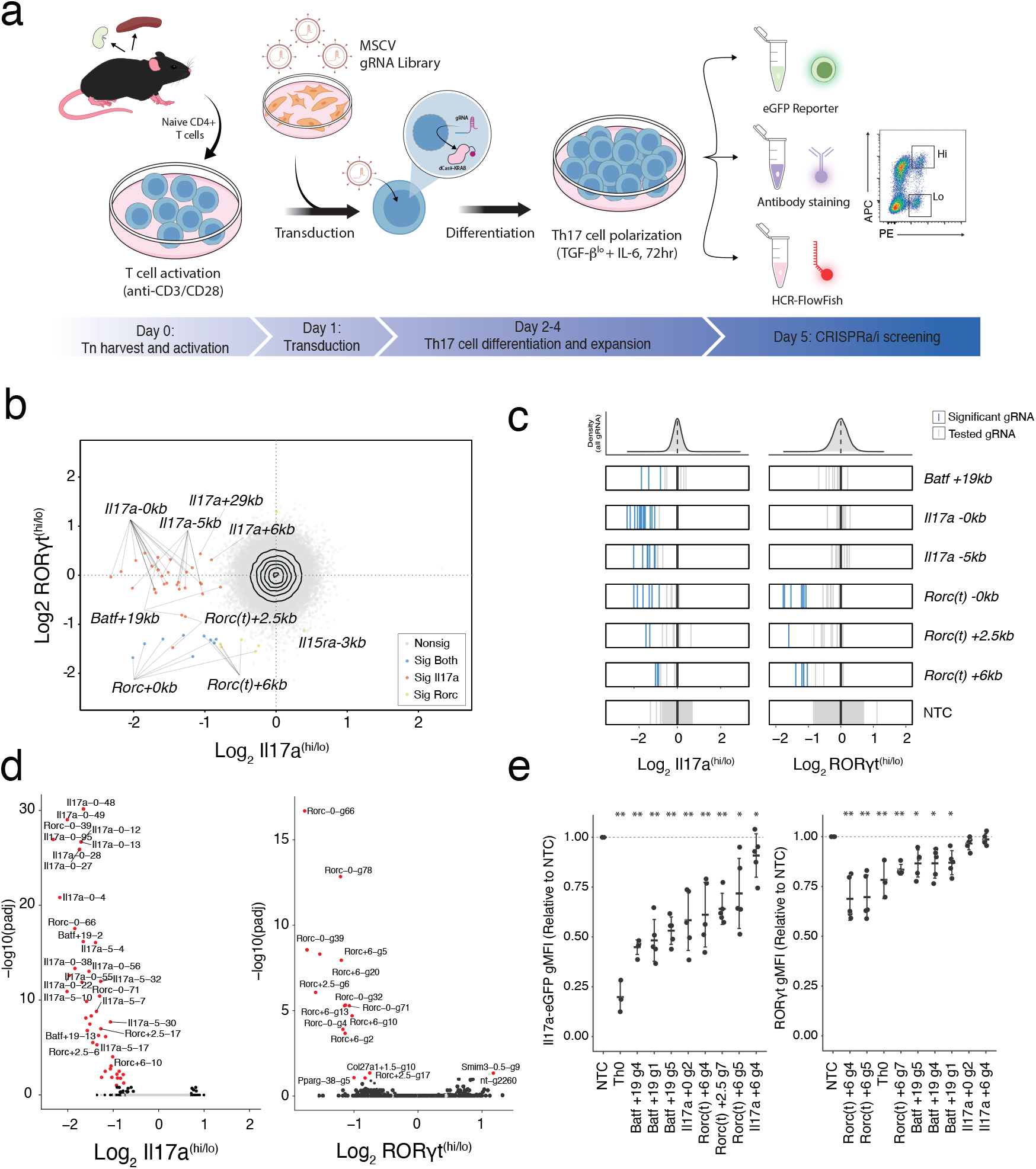
CRISPRi screening identifies functional regulatory elements controlling Th17 cell fate. **a)** Schematic of the CRISPR-based screening workflow for identifying regulatory elements involved in Th17 cell differentiation and gene regulation. Day 0: Naive CD4+ T cells were purified from spleens and lymph nodes of dCas9-KRAB mice and activated in vitro under Th0 conditions for 24h. Day 1: Cells were transduced with a retroviral library of gRNAs that targeting OCRs. Day 2: Cells were polarized under Th17 cell conditions and expanded for 72hrs. Day 4: Transduced cells were prepared for FACS using one of three readouts: i) eGFP expression (i.e. Il17a-eGFP), ii) fixed intracellular staining (i.e. RORγt, BATF), or iii) HCR-FlowFish (i.e. Il17a, Il17f). Cells were finally sorted into high or low expression bins where gRNA abundance was compared with high-throughput sequencing. **b)** Scatter plot comparing sgRNA effect sizes (Log2 fold change hi/lo) between Th17 cell differentiation noncoding CRISPRi screens with Il17a-eGFP and RORγt readouts (blue = sgRNA significant in both; padj < 0.05). **c)** Distribution of element-wise effect sizes for gRNA (vertical lines) targeting OCRs in the Il17a- and RORγt-CRISPRi screens (lines = element-targeting gRNA, blue = padj < 0.05). **d)** Volcano plots depicting sgRNA effect sizes (Log2 fold change) comparing high/low bins for Il17a-eGFP (left) and RORγt(right), with top gRNA labelled (red = padj < 0.05). **e)** Mean fluorescence intensity (MFI) of Il17a-eGFP (left) or RORγt (right) from in vitro derived Th17 cells following CRISPRi-mediated repression with individual candidate gRNAs, shown relative to non-targeting control (NTC). Box plots summarize n=5 per targeting gRNA, n=3 for Th0, n=3 for NTC. Statistical analysis was performed using one-way ANOVA with Dunnett’s test versus the NTC and sandwich standard error. Data are shown as mean ± s.e.m. relative to the NTC; * p < 0.01; ** p < 0.001

We identified 46 sgRNAs that significantly impacted Il17a production (FDR < 0.05), mapping to eight accessible regions within the *Batf*, *Rorc* and *Il17a* loci (Fig. 2b,c; Supplementary Table 4). To distinguish between regulatory elements governing Th17 cell specification versus those regulating *Il17a* expression, we repeated the screen using the Th17 cell defining transcription factor RORγt as a readout, and found the same essential regulators (Fig. 2b,d) Conversely, sgRNAs targeting the *Il17a* locus had no effect on RORγt levels, validating they regulate cytokine production downstream of subset specification. Thus, our screens identify three regulatory elements (*Batf +19kb, Rorc(t) +2.5kb, and Rorc(t) +6kb*) that drive Th17 cell polarization and two (*Il17a -5kb, and Il17a +29kb*) that specifically regulate cytokine production (Fig. 2e).

Although thousands of regions gain accessibility during polarization, our genome-wide CRISPRi approach revealed that only a small core set is individually sufficient to alter Th17 cell differentiation. This functional sparsity suggests that the Th17 cell regulatory program is both highly buffered against perturbation and dependent on a handful of subset-defining OCRs to stabilize subset identity. Having identified these essential elements at the *Batf*, *Rorc* and *Il17a* loci, we next sought to resolve the local *cis*-regulatory architecture controlling each of these genes. To achieve this, we performed complementary CRISPR-based screens targeting all Th17 cell OCRs within at least 1 Mb of each target TSS using gene-specific transcriptional or protein readouts to map local regulatory function in greater detail.

### Shared regulatory elements control *Il17a* and *Il17f* expression

We began with the *Il17* locus, where the adjacent *Il17a* and *Il17f* genes encode the core Th17 cell effector cytokines. We screened 83 OCRs across the locus using CRISPRi (Supplementary Table 3), and simultaneously quantified *Il17a* and *Il17f* mRNA abundance using multiplexed HCR- FlowFISH^34^ to directly capture transcriptional dynamics (Fig. 2a). We revealed 15 functional OCRs controlling *Il17a* and/or *Il17f* expression, including both promoters and nine distal enhancers within 50kbp (Supplementary Table 4). Notably, we found that perturbation of the *Il17a -5kb* and *+29kb* distal enhancers - both previously detected in our differentiation screen and reported by others^35,36^ - resulted in reduced expression of both Il17a and Il17f (Fig. 3a-c), suggesting a hub-like regulatory property. These dual-acting enhancers were further distinguished by element- specific differences in regulatory effect. Perturbation of the *Il17a +29kb* element strongly repressed *Il17f* but only modestly reduced *Il17a*, revealing asymmetric regulatory control, whereas targeting the *Il17a -5kb* element comparably repressed both *Il17a* and *Il17f* (Fig. 3c-f). Beyond these shared enhancers, we identified a broader network of gene-specific regulators including three *Il17a*-exclusive and four *Il17f-*exclusive functional OCRs, implicating both shared and specific *cis*-regulatory architecture at this locus (Fig. 3c-f).

**Figure 3.**
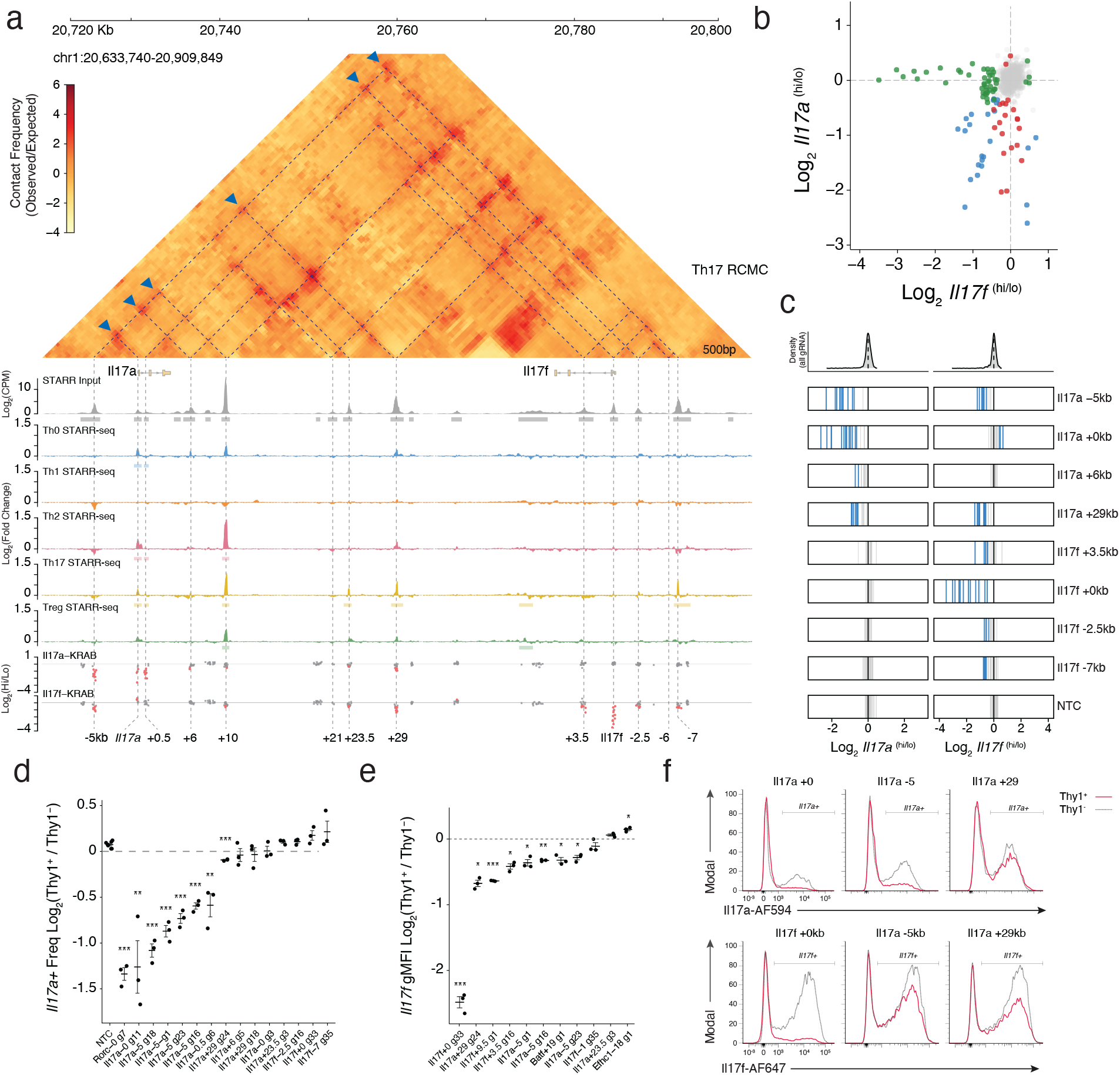
Targeted functional screen reveals dual-acting regulators of Il17a and Il17f. **a)** Genome browser plot of the Il17a / Il17f locus (70kb window) integrating 500bp resolution region capture Micro-C (RCMC; ICE balanced, normalized by observed/expected), with 3D contacts annotated by dashed line and Il17a-5 enhancer contacts indicated by blue triangles; ATAC-STARR-seq pooled input DNA library coverage track containing DNA fragments from Th0, Th1, Th2, Th17, and Treg ATAC-seq (grey); ATAC-STARR-seq activity score (Log2 fold change CPM) from Th0 (blue), Th1 (orange), Th2 (red), Th17 (yellow) and Treg (green) RNA versus Input DNA; Effect sizes for gRNA in CRISPRi for Il17a and Il17f (grey = tested; red = FDR < 0.05). OCRs are labeled with direction (+/-) and distance (in Kbp) relative to nearest gene. **b)** Scatter plot comparing sgRNA effect sizes (Log2 fold change high vs low bin) for CRISPRi screens using Il17a and Il17f reporters (green = only Il17f, red = only Il17a, blue = both, grey = non-significant; FDR < 0.05). **c)** Distribution of elementwise sgRNA effect sizes grouped by top functional OCRs in both Il17a (left) and Il17f (right) CRISPRi screens (lines = tested gRNA per element, blue = FDR < 0.05). Density plot (top) shows distribution of effect sizes for all gRNA. **d)** Flow cytometry analysis summarizing frequency of IL-17a+ cells or **e)** geometric MFI of Il17f (HCR-FlowFish) expression from in vitro derived Th17 cells following CRISPRi-mediated perturbation with candidate gRNAs. **f)** Representative stacked histograms to show distribution of in vitro derived Th17 cell Il17a and Il17f signal (red) relative to non-transduced (grey) following CRISPRi-mediated repression with top candidate single gRNA. Statistical analysis was performed using one-way ANOVA with Dunnett’s post- hoc test versus NTC and sandwich standard error **(d)** or one-sample t-tests with Benjamini-Hochberg correction **(e)**. Data are shown as mean ± s.e.m. for gRNA-transduced (Thy1.1+) relative to non-transduced (Thy1.1-) cell signal; *** p< 0.001; ** p < 0.0001; * p<0.05.

To determine how these elements engage with their target promoters, we mapped the 3D chromatin topology of the locus using Region Capture Micro-C (RCMC) for in vitro-derived Th17 cells. At 500 bp resolution, we resolved multiple interconnected subdomains demarcated by pronounced focal contacts at *Il17a* -5kb, +10kb, +29kb and *Il17f -7kb*. Crucially, this physical architecture was absent in Tn CD4+ T cells and lacked CTCF ChIP-seq signal (Supplementary Fig. 7a), indicating that interactions across the full locus were established *de novo* during Th17 cell polarization through *cis*-regulatory signalling. This 3D topology now provides the structural basis to explain the multi-enhancer regulatory logic we observed using CRISPRi.

Specifically, we found that functional OCRs determined by CRISPRi display high concordance with 3D contacts, with almost all forming physical interactions between each other (Fig. 3a). However, many functional elements at this locus do not engage a gene-promoter directly and instead interact exclusively via enhancer-enhancer (E-E) intermediates. For instance, the *Il17a - 5kb* element forms the primary physical contact with the *Il17a* promoter while simultaneously interacting with five other distal regions (Fig. 3a; blue triangles). Importantly, these intermediate OCRs are enriched for regulatory potential, as measured by ATAC-STARR-seq, yet lack direct promoter connectivity in their endogenous genomic context. Therefore, these data support a model where the *Il17a -5kb* enhancer acts as a locus control hub, likely integrating inputs from distal elements to then relay an aggregate regulatory signal to the promoter^34^.

### Novel Gene Regulatory Elements Controlling RORγt Expression

We next applied our targeted functional screening approach to the Th17 cell master regulator, *Rorc* (Fig. 2a). To define the endogenous regulatory landscape at this locus, we performed parallel CRISPRi and CRISPRa screens in Th17 cells targeting all 80 OCRs within a 2 Mbp window of the *Rorc(t)* transcriptional start site (Supplementary Table 3). This dual modality approach enabled us to define elements required for maintaining *Rorc(t)* expression, and those sufficient to increase its activity. Using RORγt protein levels as a direct readout of regulatory activity, we compared sgRNA enrichment between RORγt-high and -low populations to quantify enhancer function.

We identified seven OCRs that significantly regulated RORγt protein levels in CRISPR screening (Supplementary Table 4). Building on our primary Th17 cell differentiation screen (Fig. 2) and established regulatory maps^35–37^, our locus-specific CRISPRi approach independently converged on all known Rorc(t) regulators while revealing a previously undescribed *Rorc(t) +2.5kb* functional enhancer (Fig. 4a-c). Notably, and in contrast to the expansive network of the *Il17* locus, most of the *Rorc(t)* functional elements were constrained to a narrow 14kb window flanking the promoter (Fig 4a,d). Beyond this core cluster, most distal OCRs showed no detectable single-element effects on *Rorc(t)* expression in CRISPRi, suggesting a compact regulatory network controls this locus.

**Figure 4.**
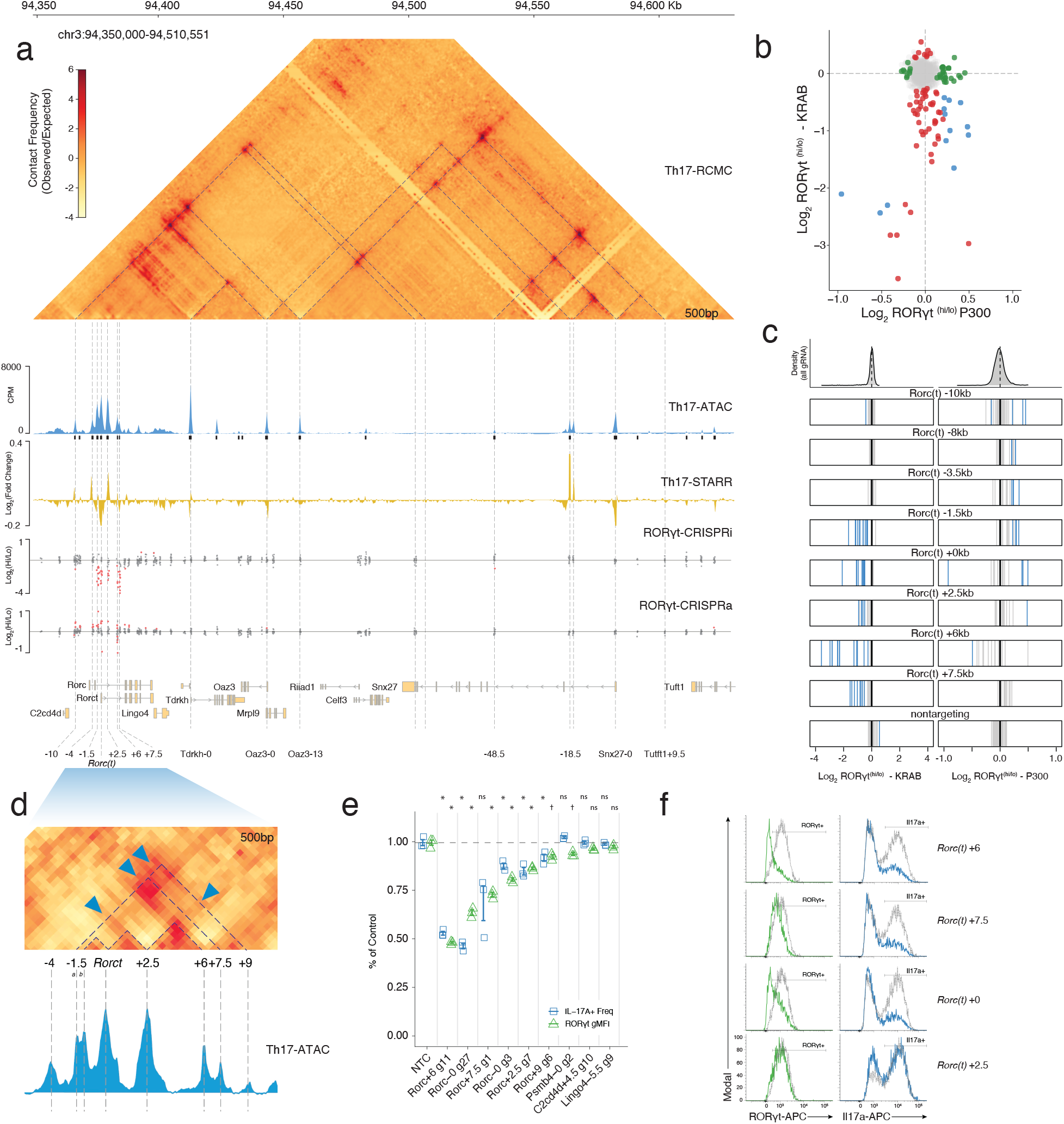
Dual effector CRISPRi and CRISPRa screen of RORγt reveal concentrated network of regulatory elements. **a)** Genome browser view of Rorc and surrounding region (200k bp region) integrating 500bp RCMC contact map (ICE balanced, observed/expected normalization) and 3D interactions annotated by dotted lines; ATAC-STARR pooled Input library (blue), Th17 cell ATAC-STARR-seq activity score (Log2 CPM (RNA / DNA); yellow); CRISPRi- and CRISPRa-RORγt effect size (red = sgRNA FDR < 0.05; grey = tested); and OCR annotations label the direction (+/-) and distance (in Kbp) from nearest gene. **b)** Scatter plot comparing sgRNA effect sizes (Log2 fold change RORγt high vs low bins) from CRISPRi and CRISPRa screens (green = CRISPRa only; red = CRISPRi only; blue = both; grey = nonsignificant; FDR < 0.05). **c)** Element-wise distribution of effect sizes for both CRISPRi (left) and CRISPRa (right) (lines = element-tested gRNA; blue = FDR < 0.05) **d)** Zoomed in RCMC contact map (500bp resolution) focusing on the proximal RORγt locus (16kb window) with 3D contacts annotated as dotted lines, notable contact enrichments labelled with blue triangles, and corresponding Th17 cell ATAC-seq coverage track (blue) **e)** Frequency of IL-17a (blue) or MFI of RORγt (green) relative to non-targeting control (NTC) for in vitro derived Th17 cells following CRISPRi-mediated perturbation with top candidate gRNA from RORγt screening. **f)** Representative stacked histograms depicting RORγt and IL17a flow cytometry signal for Th17 cells transduced (Thy1+; blue/green) or nontransduced (Thy1-; grey) with candidate gRNAs. Statistical analysis was performed using one-way ANOVA with Dunnett’s test versus the NTC and sandwich standard error. Data are shown as mean ±

Within that *Rorc(t)* regulatory core (Fig. 4d), CRISPRi-mediated repression and CRISPRa-mediated activation of the immediate promoter-proximal OCRs had symmetrical, opposing effects on RORγt protein levels, revealing a dual-regulatory capacity (Fig. 4c). However, the regulatory function of the promoter-distal OCRs was segregated by CRISPR modality. The downstream functional OCRs (+2.5 +6, +7.5) were necessary for the regulation of RORγt expression, with endogenous perturbations resulting in strong repressive effects, yet they were largely unresponsive to further activation (Fig 4a-c). Strikingly, targeting CRISPRa to the +6kb downstream element also resulted in repression of *Rorc(t)*, rather than activation (Fig. 4b,c). This paradoxical effect is not well understood, but suggests that exogenous recruitment of bulky protein complexes may be sufficient to sterically hinder regulatory function in specific instances where regulation relies on precise biophysical interactions^2^.

Conversely, OCRs upstream of *Rorc(t)* (−4 kb to -10 kb) were dispensable in the repression screen but drove increased RORγt expression following CRISPRa (Fig. 4a-c). The functional divergence between these sets of OCRs aligned with their intrinsic regulatory potential measured by ATAC-STARR-seq (Supplementary Fig 6c). Episomal activity was relatively robust at many of the CRISPRa-responsive OCRs, but weak at most of those affected by CRISPRi (Fig. 4a; Supplementary Fig. 6c). A notable exception was at the +2.5kb OCR, where CRISPRi-mediated repression and STARR-seq activity signal converged, suggesting sequence and endogenous chromatin context contribute to the regulatory function of *Rorc(t) +2.5kb* in *Rorc(t)* expression. Targeting these core elements with CRISPRi in Th17 cells individually confirmed that single perturbations resulted in reductions to RORγt and its downstream target IL-17A (Fig. 4e-f). These effects varied between individual gRNA within a given element (Fig. 4e-f).

We then mapped the 3D physical interactions of functional regions within the *Rorc(t)* locus using region-capture micro-C in Th17 cells (Fig. 4a,d). At 500-bp resolution we found the broader locus to be organized by multiple prominent subdomains, whereas the promoter-proximal regulatory core was resolved into nested micro-domains that largely correspond with functional OCRs (Fig 4d). By comparing these 3D chromatin contacts to those in Tn cells, we determined that the distal architecture of the locus was pre-established, but the specific micro-topology associated with the *Rorc(t)* promoter was formed de novo upon Th17 cell differentiation, and without an associated CTCF ChIP-seq signal(Supplementary Fig. 7b). By overlaying our functional screening data onto this high-resolution contact map, we observed distinct connectivity between the core *Rorc(t)* OCRs (Fig. 4a). Promoter-proximal functional OCRs (−1.5 kb and +2.5 kb) form direct physical contacts with the TSS, while the distal downstream elements (+6 kb and +7.5 kb) interact indirectly, and only with these proximal enhancers (Fig. 4d). Specifically, the +6 kb element contacts the +2.5 kb region, and both distal elements interact with the -1.5 kb enhancer. These data suggest a model in which the downstream functional OCRs are tethered to the TSS through proximal enhancers, forming functionally necessary, hub-like, enhancer-enhancer interactions. The contrast between strong regulatory signal in CRISPRi and low regulatory potential in STARR-seq at these distal regions points to a structural rather than directly regulatory role, possibly stabilizing the regulatory micro-domain through multi-enhancer connectivity.

### A novel intronic enhancer at *Batf +*19kb controls *Batf* expression and drives Th17 cell polarization

Finally, to identify regulatory elements controlling expression of *Batf,* an essential regulator of Th17 cell differentiation, we applied the dual-modality CRISPR screening approach described above, targeting 114 OCRs within 1 Mbp of the *Batf* promoter (Supplementary Table 3). We quantified regulatory effects by directly measuring BATF protein levels via intracellular staining and flow cytometry of in vitro-derived Th17 cells and compared sgRNA enrichments between BATF-high versus -low cell populations (Fig 2a).

Compared to the *Il17a/f* and *Rorc* loci, *Batf* featured a comparably sparse regulatory landscape, with a more direct regulatory logic (Fig. 5a). Using CRISPRi, we detected regulatory function at two promoter-proximal (*Batf +0kb, +0.5kb*) and two distal OCRs (*Batf* +19kb, *Flvcr2* -16.5kb). We further identified nine CRISPRa-responsive regions within a 100kb window of the promoter (Fig. 5a-c; Supplementary Table 4). Like *Rorc*, the *Batf* promoter-proximal elements were sensitive to both repression and activation (Fig. 5b,c). However, at this locus, most CRISPRa-responsive regions were dispensable in CRISPRi and had low Th17 cell STARR-seq signal, suggesting they may be poised or context-dependent elements that lack the necessary cofactors for endogenous function in Th17 cells. Instead, the distal *Batf +19kb* and *Flvcr2 -16.5kb* OCRs displayed both CRISPRi necessity and STARR-seq intrinsic activity, consistent with the functional convergence we observed at the Il17a/f locus and the *Rorc(t) +2.5kb* hub (Fig. 5a-c; Supplementary Figure 6b- d). Collectively, these data indicate that *Batf* expression is primarily driven by two high potency distal enhancers.

**Figure 5.**
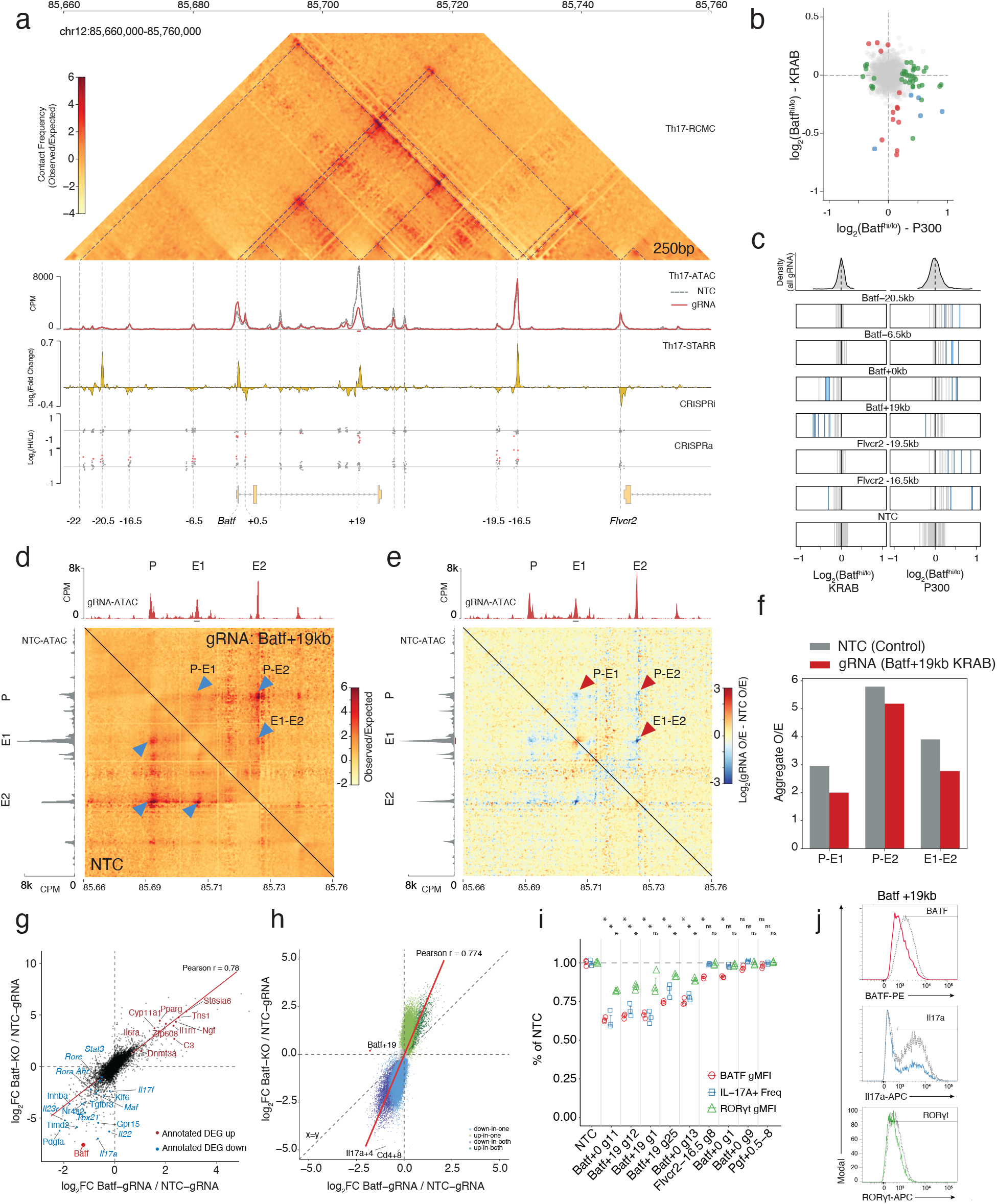
The +19kb Batf enhancer is a structural and functional core regulatory element in Th17 cells. **a)** Multimodal view of the Batf locus (100k bp window). Top: Region-capture Micro-C (RCMC) contact map (250bp resolution; ICE balanced), with interactions annotated by dotted lines. Tracks display Th17 cell ATAC-seq coverage by condition (non-targeting control [NTC] = grey; +19kb CRISPRi = red), Th17 cell ATAC-STARR-seq activity (Log2 CPM RNA / DNA; yellow), and CRISPRi/CRISPRa screen effect sizes (points indicate tested sgRNA, red = FDR < 0.05). Enhancers are annotated by distance (kb) and direction (+/-) relative to the Batf TSS. **b)** Scatter plot comparing CRISPRi versus CRISPRa effect sizes (Log2 fold change) for all tested sgRNA. Points coloured by significance (FDR < 0.05). **c)** Distribution of sgRNA effect sizes at selected elements from CRISPRi (left) and CRISPRa (right) screens (blue = significant; grey = tested) **d)** Comparison of RCMC contact frequency (500bp resolution) at the Batf locus following transduction with Batf +19kb-targeting (top) or NTC (bottom) sgRNAs in dCas9-KRAB Th17 cells. **e)** Differential contact map showing Log2 fold-change in interaction frequency (Batf +19kb sgRNA / NTC) **f)** Aggregate Peak Analysis quantifying contact frequency of interactions between the Batf-TSS (P), Batf +19kb (E1) and Batf +43kb (E2) elements in CRISPRi-mediated Batf +19kb perturbed Th17 cells (red) versus NTC (grey). **g)** Quantitative comparison of transcriptomic changes measured by RNA-seq (Log2 fold-changes relative to control) or **h)** chromatin accessibility changes by ATAC-seq (Log2 fold-change relative to control) in Batf-/- (BATF-KO) and CRISPRi-mediated Batf +19kb enhancer perturbation (Batf-gRNA) of in vitro derived Th17 cells (RNA Pearson’s r = 0.78; ATAC Pearson’s r = 0.774). i) MFI of BATF (red) or RORγt (green) , and frequency of IL-17A+ (blue) from in vitro derived Th17 cells following CRISPRi-mediated repression of candidate OCRs with single gRNA relative to non-targeting control. Box plots summarise n=3 biological replicates **j)** Representative stacked histograms for BATF (red) IL-17a (blue) and RORγt (green) protein levels in Th17 cells following CRISPRi-mediated repression of Batf +19kb enhancer compared to nontargeting control (grey). Statistical analysis was performed using one-way ANOVA with Dunnett’s test versus the NTC and sandwich standard errors. Data are shown as mean ± s.e.m. relative to the NTC; * p < 0.001.

To determine whether these distal enhancers physical contact the *Batf* promoter, we performed region-capture Micro-C on in vitro-derived Th17 cells (Fig. 5a). At 250bp resolution, we resolved multiple contact domains spanning 80kb across the locus, most of which are pre-established in the naive T cell (Fig. 5a; Supplementary Fig. 7c). Upstream of *Batf*, we identified diffuse contacts forming faint subdomains that encompass many CRISPRa-responsive elements (Fig. 5a), further supporting their possible poised function for an alternative cell state. In contrast, the chromatin topology downstream of *Batf* was structurally pronounced (Fig 5a). Distinct from the topological complexity observed at *Il17a/f* and *Rorc*, the distal *Batf* CRISPR-responsive elements form simple, direct loops with the promoter. Central to this organization, the *Batf +19kb* OCR anchors two nested subdomains within a larger structure, connecting both to the promoter and to downstream distal elements (Fig. 5a).

Given the essential role of the *Batf +19kb* OCR in Th17 cell polarization, we decided to investigate whether its activity was required to maintain its nested 3D chromatin architecture. Upon targeted repression of *Batf +19kb* with dCas9-KRAB we observed specific reductions of the internal chromatin contacts, depleting both enhancer-promoter and enhancer-enhancer loops (Fig. 5d,e). Aggregate peak analysis (APA) confirmed a 1.5-fold depletion in contact frequency for interactions between the *Batf +19kb* element and promoter (E-P) and *Batf +19kb* and *Flvcr2 -16.5kb* (E-E; Fig. 5f). Remarkably, the topology of the broader locus remained intact, with minimal reduction to contacts between the promoter and the subdomain boundary at *Flvcr2 -16.5kb* (Fig. 5f). Taken together, we show that the selective disruption of enhancer-promoter loops leads to a reduction of BATF protein levels that was sufficient to disrupt Th17 cell phenotypes (Fig. 5i,j).

To quantify the extent of this enhancer-mediated regulatory control, we measured the genome-wide effects of perturbing the *Batf +19kb* enhancer in Th17 cells using RNA- and ATAC-seq. Relative to control, enhancer repression induced thousands of changes in gene expression (Supplementary Table 5) and chromatin accessibility (Supplementary Table 6). We then compared the magnitude of these changes to those observed in cells from *Batf^−/−^* mice^24^ against control. This analysis revealed a striking correlation between the effect sizes of *Batf +19kb* enhancer repression, and those from *Batf^-/-^* for both gene expression (Fig. 5g; Pearson r = 0.74 ) and accessibility (Fig. 5h; r = 0.78). Core Th17 cell genes (e.g., *Rorc*, *Stat3*, *Il17a, Il23r*) and known *Batf* targets were downregulated with comparable magnitudes in both enhancer-CRISPRi and gene-KO groups, while metabolic and epigenetic regulators were similarly upregulated (Fig. 5g).

Although the magnitude of change in chromatin accessibility was much stronger in the Batf^-/-^ condition, the direction of change was overwhelmingly concordant (Fig. 5h). Nevertheless, a proportion of *Batf^-/-^* specific differential OCRs were determined as not significantly altered following CRISPRi repression of the *Batf +19kb* enhancer. Peaks losing accessibility exclusively in Batf^-/-^ cells were enriched for RUNX and nuclear receptor (NR) motifs consistent with RORγt binding sites, while peaks affected in both conditions were enriched for BACH and NFAT (Supplementary Fig. 8). Together, these data demonstrate that repressing the *Batf +19kb* enhancer alone was sufficient to closely recapitulate the genomic effects of a full knockout, while condition specific motif differences suggest distinct pioneering and maintenance roles of BATF these elements.

### In vivo validation of essential Th17 cell regulatory elements

While the differentiation models used above mimic *in vivo* signalling pathways, many aspects of T cell subset specification cannot be modeled *in vitro*. Therefore, we validated our top functional regulatory elements *in vivo* using the 7B8 TCR transgenic model^38^ crossed with our dCas9-KRAB mouse line. T cells from this system recognize the segmented filamentous bacteria (SFB), enabling antigen specific Th17 cell polarization in the gut microenvironment. To evaluate enhancer essentiality in this physiological context, we delivered sgRNAs to the CD4+ T cells from 7B8-KRAB mice to repress the distal OCRs *Il17a -5kb*, *Rorc(t) +2.5kb*, and *Batf +19kb*, as well as the *Rorc(t)* promoter-proximal OCR *Rorc(t) +0kb*. After adoptive transfer and SFB colonization, we analyzed donor T cell recovery and phenotypes from the small intestine lamina propria (siLP) and compared them to non-targeting controls (Fig. 6a).

**Figure 6.**
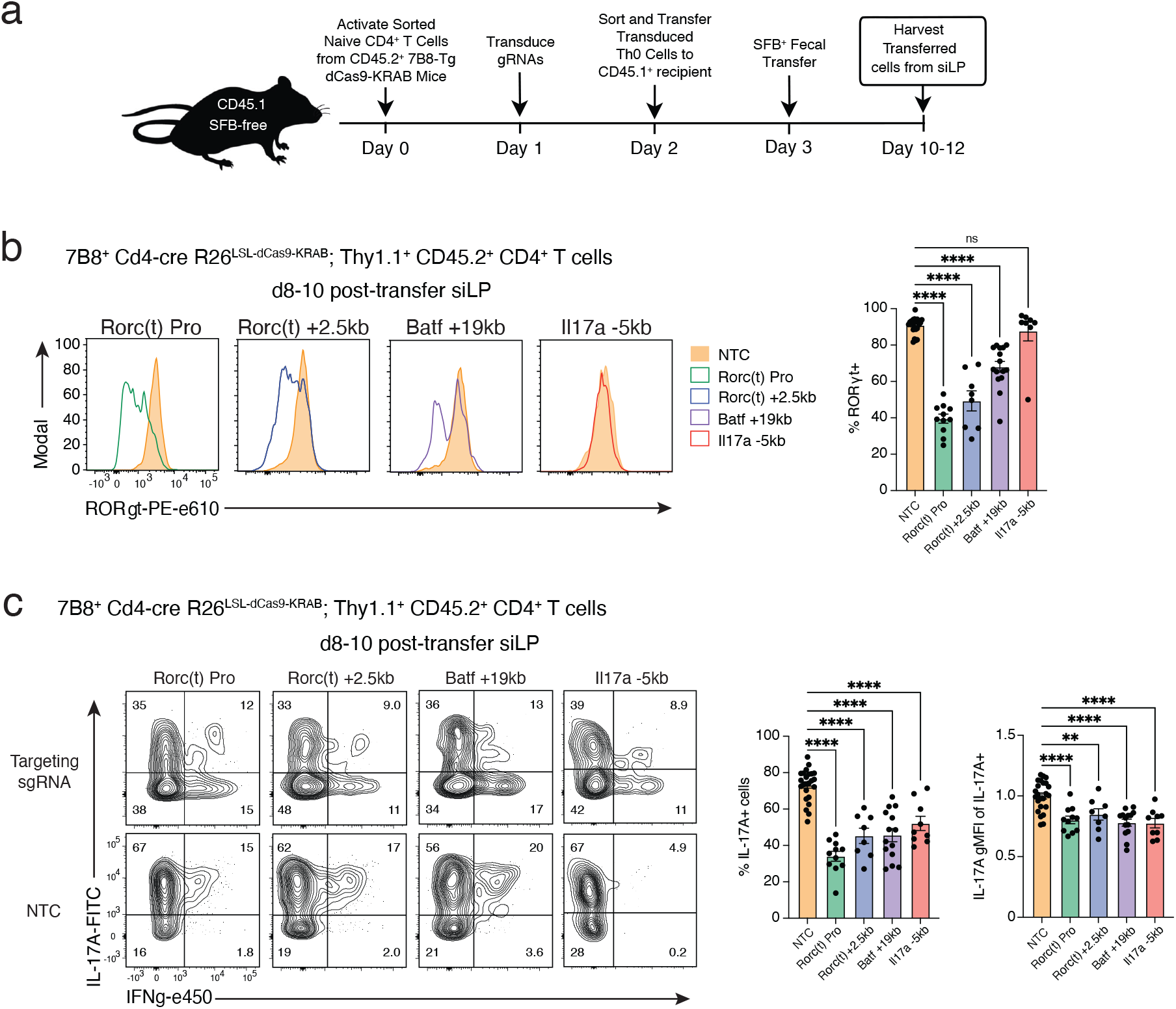
In vivo validation of Th17 cell enhancers in the 7B8 adoptive transfer model with SFB-driven differentiation. **a)** Schematic of the experimental workflow for in vivo validation of sgRNAs using the 7B8-KRAB transfer and SFB colonization system. **b-c)**, Flow cytometry analysis of donor Thy1.1+ CD45.2+ CD4+ T cells derived from 7B8+ Cd4-cre Rosa26LSL-dCas9-KRAB mice transduced with non-targeting control (NTC), Rorc(t) promoter (Rorc(t) Pro), Rorc(t) +2.5 kb, Batf +19 kb, or Il17a −5 kb sgRNAs and recovered from small intestinal lamina propria (siLP) of CD45.1 recipient mice 8-10 days after transfer. Columns denote the targeted regulatory element (n = 8-24; two to seven independent experiments per sgRNA condition). **b)** Representative RORγt histograms for sgRNA-targeted elements and experiment-matched NTC donor cells. Summary plots quantify the frequency of RORγt+ cells within the donor population following repression of the indicated enhancer elements. **c)** Representative intracellular cytokine staining for IL-17A and IFNγ in sgRNA-targeted and experiment-matched NTC donor cells. Summary plots show the frequency of IL-17A producing donor cells and the geometric mean fluorescence intensity (gMFI) of IL-17A within the IL-17A+ population. Donor Thy1.1+ CD45.2+ CD4+ T cells were gated as CD4+ Foxp3- CD3ε+ TCRβ+ CD45.2+ Thy1.1+. Cytokine expression in c was measured after 4 h PMA/ionomycin stimulation. Statistical analysis was performed using one-way ANOVA and Dunnett’s test to the NTC **(b-c)**. Data are shown as mean ± s.e.m.; **P < 0.01; ****P < 0.0001. Numbers in flow plots indicate the percentage of cells within the indicated gates.

For all perturbation groups, we successfully recovered in vivo differentiated donor 7B8+ Th17 cells from the siLP (Supplementary Fig. 9a). Consistent with our in vitro findings, targeted repression of functional OCRs at lineage defining loci (*Rorc(t) +0, Rorc(t) +2.5kb, Batf +19kb*) significantly impaired Th17 cell specification. *In vivo* CRISPRi at these loci resulted in decreased proportions of RORγt+ cells between 1.3- to 2.3-fold (Fig. 6b), and reduced IL-17a+ cell populations between 1.6- to 2.2-fold (Fig. 6c). In contrast, our effector-specific perturbation (*Il17a -5kb*) selectively disrupted IL17a production, reducing IL17a+ cells 1.4-fold without affecting the frequency of RORγt+ cells (Fig. 6b-c). Strikingly, the dramatic down regulation of RORγt induced by targeting either the Rorc(t) promoter or the +2.5kb enhancer resulted in increased levels of IFNγ for the IFNγ-producing 7B8+ T cells (Supplementary Fig. 9d), indicating subset plasticity towards a Th1-like state, positioning these noncoding elements as important regulators of Th17 cell identity. Collectively, these data demonstrate that CRISPRi-mediated perturbation of a single distal enhancer is sufficient to profoundly shape Th17 cell specification and alter immune function in mucosal tissues.

## Discussion

This study provides an integrated framework for understanding cis-regulatory function in mouse CD4+ T cells. By decoupling sequence potential from the native chromatin context using a pooled ATAC-STARR-seq approach, we demonstrate that only a fraction of all CD4+ OCRs exhibit intrinsic regulatory activity; however, this activity remains largely consistent when compared across T cell subsets. Despite this shared active potential, we resolve the degree to which activity is biased towards individual subsets, identifying OCRs with subset-preferred activity that predominates in the effector lineages. Focusing on Th17 cell polarization, we identify a sparse set of enhancers with endogenous function in subset specification. Integrating high-resolution chromatin conformation, we show how functional elements of the *Il17*, *Rorc(t),* and *Batf* loci each assemble into nested, multi-enhancer hubs with physical interactions that explain their regulatory function. Crucially, we show that epigenetic perturbation of a single enhancer at the *Batf* locus is sufficient to disrupt 3D chromatin contacts and impair downstream cellular phenotypes, directly linking enhancer function to physical connectivity.

Reporter assays provide a broad assessment of functional activity by testing the intrinsic potential of DNA sequence in a permissive episomal context. Several features of the STARR-seq assay warrant caution in interpretation. First, the fraction of active elements that we detect may be conditioned on the choice of minimal promoter. Recent work shows promoters differ substantially in the magnitude of responsiveness to enhancer activation {tang 2026; stark 2017}. Our single promoter design could therefore constrain the apparent number of active elements, making this enhancer set a conservative estimate. However, this alone cannot account for the central observation that regulatory activity is decoupled from chromatin accessibility and its remodelling during polarization, nor the relative activity of enhancers observed between cell subsets. Second, GC- and AT-rich sequence is a known technical source of bias in genomics assays. Sequence feature analysis shows how GC and AT bias plays a major role in our negative STARR-seq signal. Although prior work indicates such assays can capture genuine repressive function^14^, we interpret repressive calls with caution.

Our subset-preferred active elements are enriched for transcription factor motifs that constitute a cell-type-consistent subset of those also enriched among the broadly shared active elements. This suggests an alternative regulatory mode where preferential activity is encoded by sequence features other than motif identity. While the underlying mechanisms remain to be understood, we hypothesize that variation in the core regulatory sequences could modulate motif affinity, thereby intrinsically tuning their response to context-specific differences in TF abundance^39,40^. Alternatively, non-additive interactions between functional motifs might render an element sensitive to specific factors present in each subset, resulting in graded rather than binary signal across cell states^41–43^. Resolving these sequence interactions will likely require base-pair resolution dissection through high-coverage synthetic libraries or sequence-models trained on dense functional data.

However, latent potential does not reliably predict endogenous function. While reporter assays define the theoretical regulatory capacity of a DNA sequence, our CRISPR screens highlight how the native chromatin context ultimately determines its functional effect. Through CRISPR-based epigenetic perturbations, we describe a core set of enhancers that control Th17 cell specification via locus-specific gene regulation. The narrow hit rate in our non-coding screens underscores both the robustness and fragility of these enhancer networks. Although many of the most dynamic Th17 cell OCRs showed no detectable single-element effect in our screens, disrupting a single regulatory hub is sufficient to induce major changes in cellular identity and function. Indeed, factors such as enhancer redundancy^41,44^, timing of perturbation, and variable gRNA efficiencies^45,46^ likely contribute to the specificity of our findings. Future studies capturing a wider range of downstream transcriptional effects may expand the functional enhancer landscape. Nevertheless, our integrated analysis converges on several core themes of gene regulation.

Our data support a model in which physical connectivity is a primary determinant of regulatory element function. While others have presented evidence for this theory^47–50^, we show how enhancer-promoter linkages are specifically related to functionally validated OCRs. In Th17 cells, the CRISPRi-responsive functional enhancers are distinguished by enhancer-enhancer physical interactions, largely without an associated CTCF loop anchor. In contrast, elements with latent activity revealed by CRISPRa do not require such connectivity but are mostly constrained to the same topological subdomains as their target gene. Moreover, the enrichment of STARR-seq sequence activity specifically at OCRs with CRISPR-validated function suggests a functional relationship between intrinsic enhancer potential, physical connectivity, and endogenous regulatory function.

The high resolution of region capture Micro-C reveals how these interactions operate in the native chromatin context. Building on prior descriptions of multivalent, hub-like enhancers^51,52^, we resolve how these elements assemble into structured regulatory networks, in which enhancer contacts are selective rather than all-to-all. Importantly, this organization is not uniform and varies across loci. For instance, at the Il17a/f locus, distal E-E interactions aggregate at the hub-like *Il17a -5kb* OCR rather than interacting with the Il17a promoter directly. At the *Rorc(t)* locus, they split between two promoter-connected regulatory hubs (+2.5kb and -1.5kb). These specific contacts form de novo during differentiation, absent in the naïve state. In contrast, the Batf locus is defined by more direct, pre-established E-P contacts. Perturbing the *Batf* +19kb hub within this preformed architecture reduces contact frequency and impairs target gene expression, demonstrating that these interactions depend on enhancer function. Together these findings have broad implications for understanding how gene regulatory networks organize distal elements to mediate gene expression during cell differentiation.

By characterizing the cis-regulatory elements of immune-cell differentiation, we find the noncoding genome to be defined by widespread functional redundancy, from which a sparse set of hub-organized enhancers emerges as essential to Th17 cell identity. The regulatory architecture measured at Th17 cell enhancer hubs begins to explain why only some enhancers are essential, advancing our understanding of the structural and functional specificity required for gene regulation. By resolving how sequence potential, enhancer function, and 3D architecture converge at essential elements, this study offers a framework for dissecting cis-regulatory logic beyond the immune system. Whether hub organization is a generalizable signature of enhancer essentiality across cell types and developmental contexts is a central question for future research.

## Supplementary Figures

**Supplementary Figure. 1.**
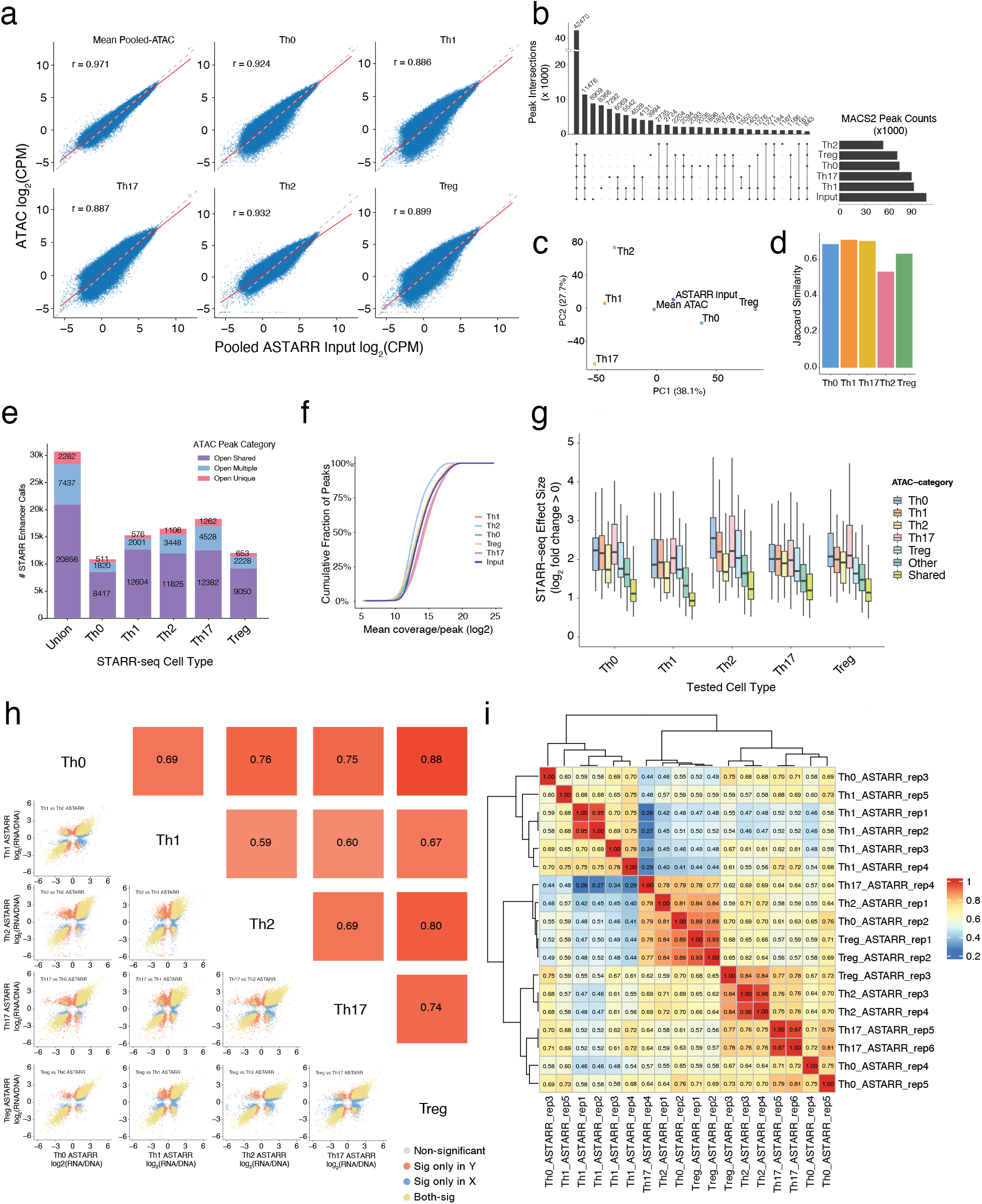
Pooled ATAC-STARR-seq library metrics. **a)** Quantitative comparison of read counts from ATAC-seq peaks for all subset-specific ATAC-seq datasets and mean count across all subsets against read counts (Log2 CPM) for the pooled ATAC-STARR-seq Input library (red line = Pearson correlation). **b)** Upset plot showing intersection of total peak calls between all ATAC-seq and STARR-seq Input libraries. Peak intersections require MACS2 FDR < 0.001 in at least 2 replicate samples. **c)** PCA of read counts (CPM) for ATAC-seq and ATAC-STARR-seq input libraries at STARR-seq tested OCRs. **d)** Jaccard index of MACS2 determined peak similarity between ATAC-seq libraries and ATAC-STARR-seq input library. **e)** ATAC-STARR-seq total enhancer calls for each subset categorized by ATAC-seq status (purple = accessible in all subsets; red = in only one subset, blue = multiple subsets). **f)** Cumulative fraction of reads in peak covering tested MACS2 peaks for ATAC and ATAC-STARR-seq input libraries. **g)** STARR-seq effect size across subsets for active peaks (FDR < 0.05, Log2FC > 0) categorized by their endogenous ATAC-seq class. **h)** Pairwise correlation plot of mean STARR-seq effect size (Log2 RNA/DNA) across replicates for all subsets tested in this study (Points = OCR, yellow = sig in both; red = sig only on Y axis; blue = sig only on X axis; grey = nonsignificant) **i)** Pairwise correlation plot of STARR-seq effect sizes (Log2 RNA/DNA) for all individual replicates.

**Supplementary Figure. 2.**
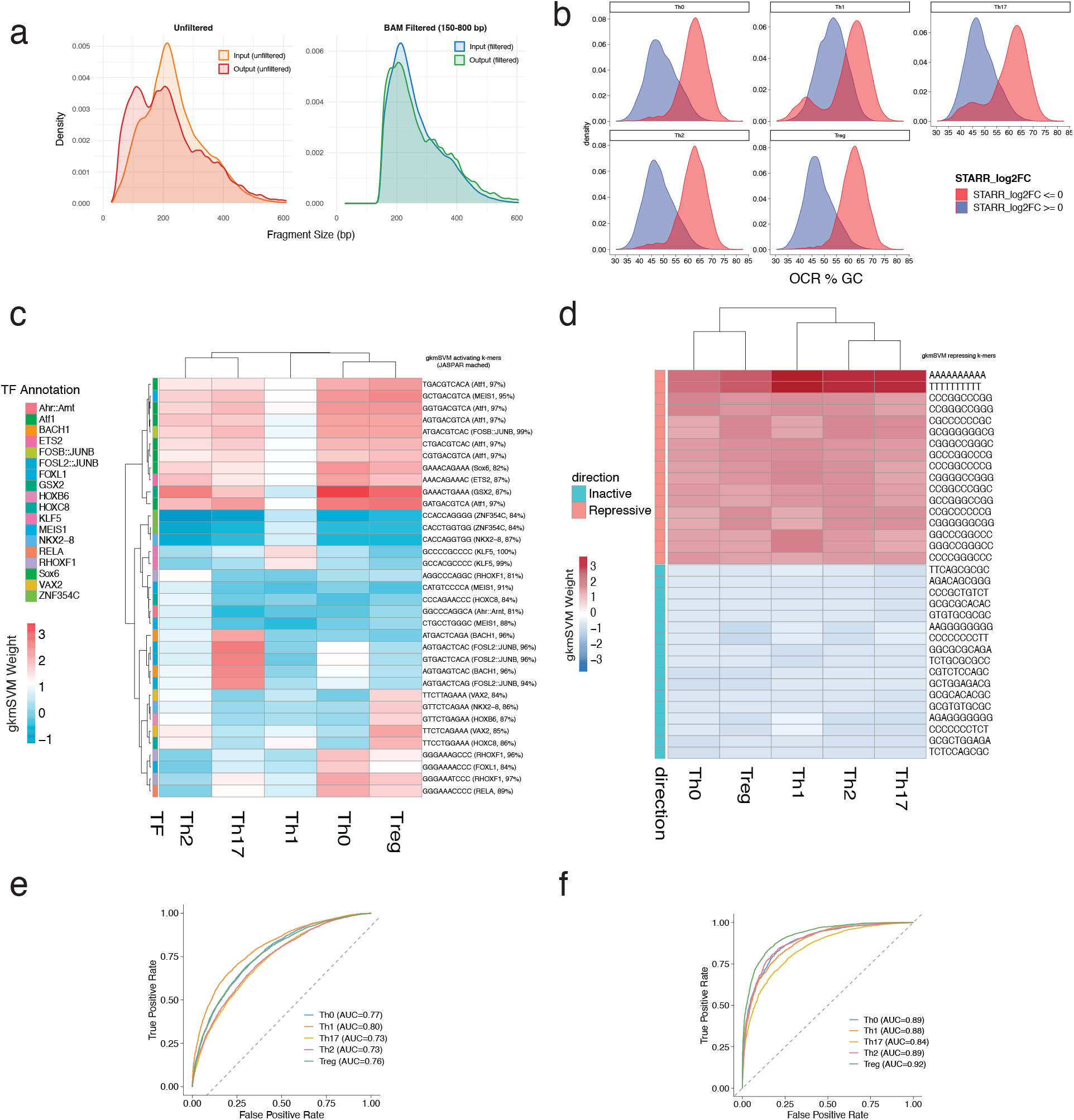
STARR seq data filtering approach. **a)** Fragment size distribution of pre-fragment filtering of ATAC-STARR-seq input libraries (orange) and RNA libraries (red), and post-filtering to keep only 150-800bp fragments input (blue) and output (green) libraries. **b)** Distribution of GC content for OCRs with positive STARR-seq signal (blue) and repressive STARR-seq signal (red). **c)** Cross-subset comparison heatmap of gkm-SVM k-mer weights from active (FDR < 0.05; Log2 fold change > 0) vs inactive ATAC-STARR OCRs and **d)** from repressive (FDR < 0.05; Log2 fold change < 0) vs inactive ATAC-STARR-seq OCRs. **e)** AUC per subset for active vs inactive gkmSVM classification. **f)** AUC per subset for repressive vs inactive gkmSVM classification

**Supplementary Figure. 3.**
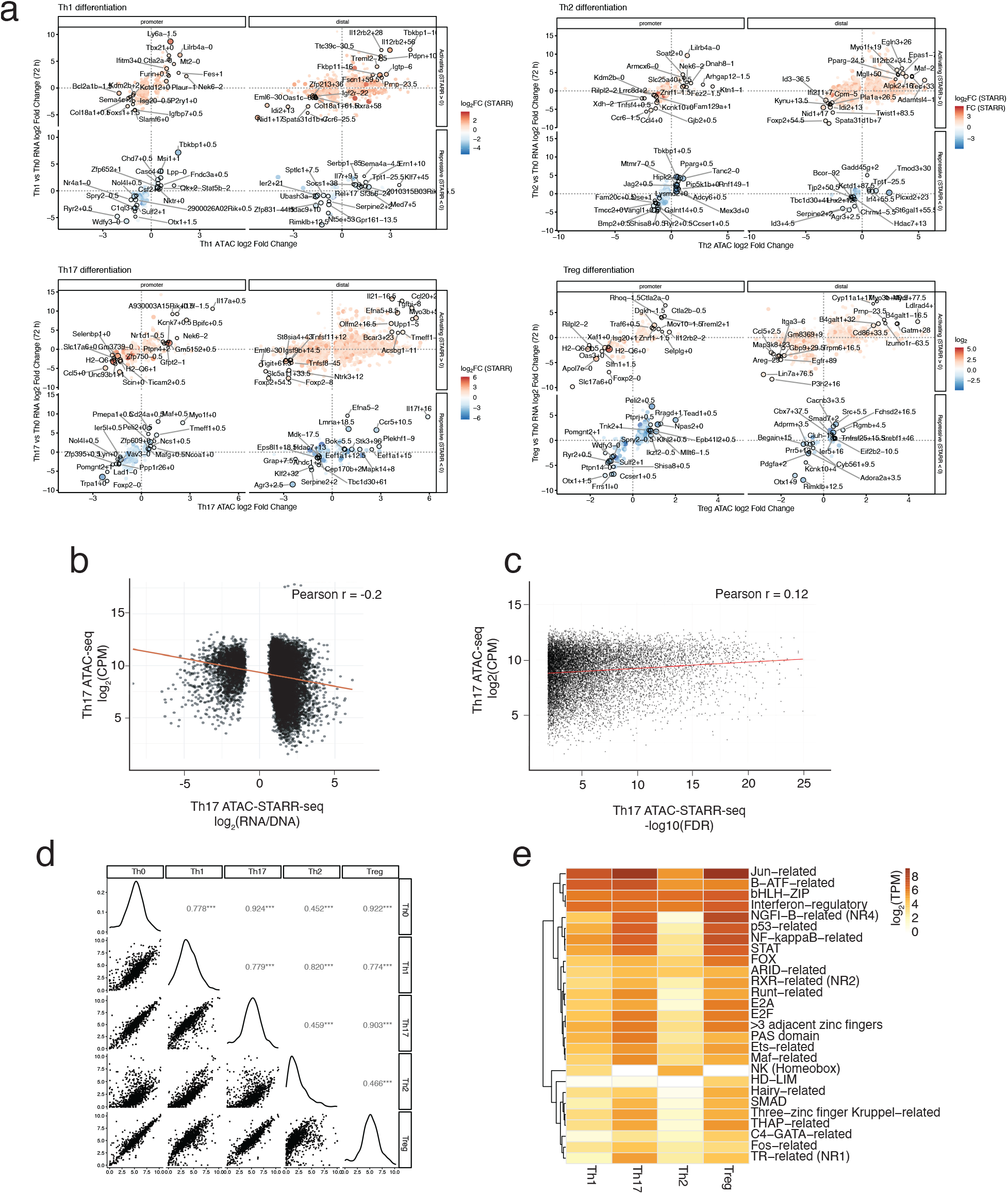
STARR-seq Correlation to Genomic and Transcriptomic Context. **a)** Correlation of differential RNA-seq (All Subsets vs Th0) and nearest differential ATAC-seq (All Subsets vs Th0) peaks, split by promoter-proximal (< 3kbp) and distal (≥ 3kbp) from TSS, activating (Log2 RNA/DNA > 0) and repressive (Log2 RNA/DNA < 0) and coloured by magnitude of STARR-seq activity (Log2 RNA/DNA). **b)** Correlation of Th17 cell ATAC-seq signal at peak regions (Log2 CPM) against Th17 cell ATAC-STARR-seq activity score for all enhancers (Pearson’s r = -0.2). **c)** Correlation of Th17 cell ATAC-seq signal at peak regions (Log2 CPM) against ATAC-STARR-seq significance (- log10(padj); Pearson’s r = 0.12) **d)** Pairwise correlation of transcript abundance (Log2 TPM) for all known transcription factors and epigenetic modifiers expressed in the CD4 T cell subsets of this study (Pearson’s correlations = 0.45-0.92). **e)** Expression levels (mean TPM) of transcription factor families shown in Fig. 1J, averaged across family members, for each CD4 T cell subset at 72hrs post polarization.

**Supplementary Figure. 4.**
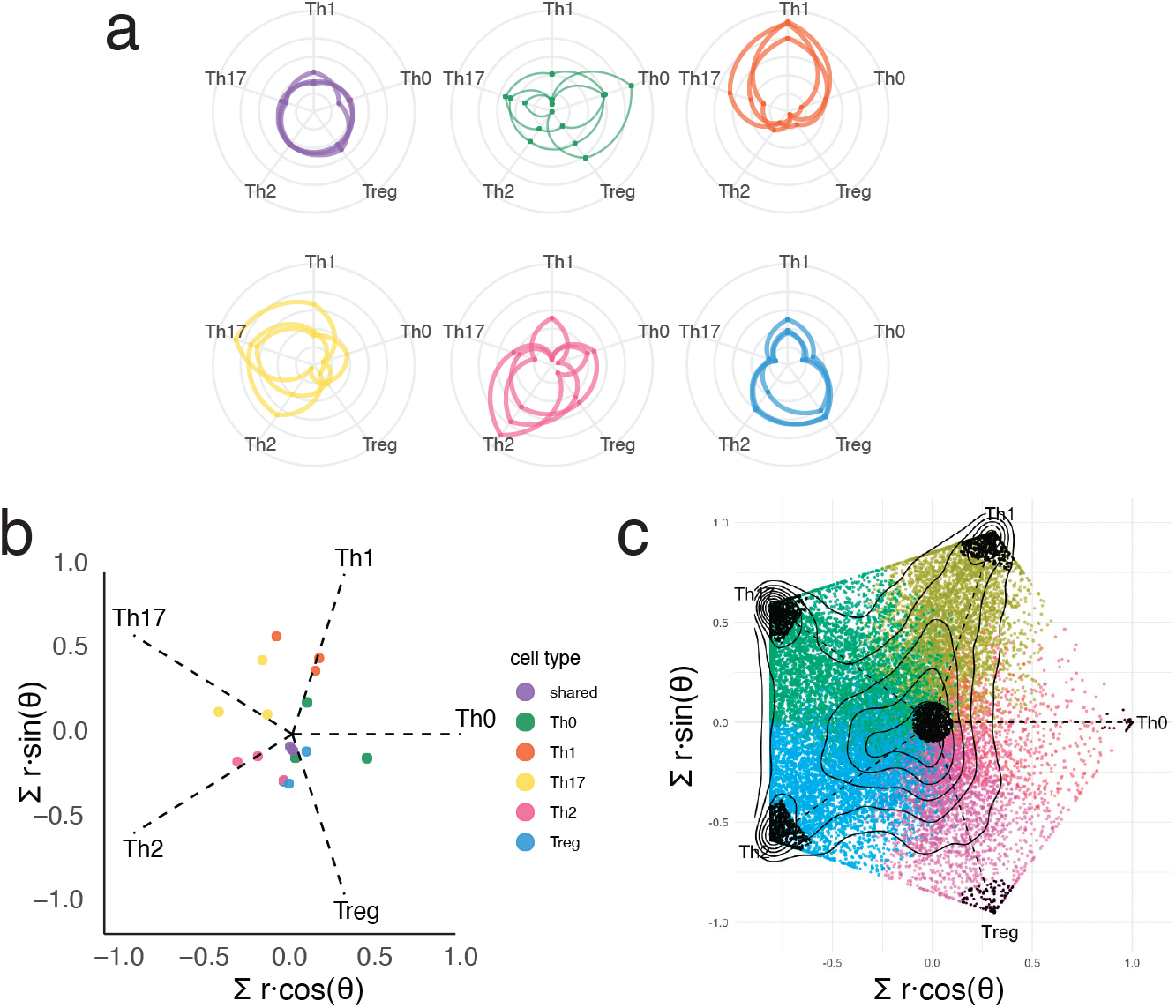
Example breakdown of radial plot projection for subset specificity analysis. **a)** Example of radial position for selected OCRs. Enhancer weighted effect size measured for each subset was plot radially along the fixed axis. Connected lines indicate a single OCR tested across all subsets. **b)** Summation of vectors for each OCR across each subset reveals the mean radial position for each OCR indicating subset preference in STARR-seq activity, and distance from center indicates the magnitude of that preferential activity. **c)** Final plot indicating the subset-preferred or subset shared enhancers used for downstream motif analysis in black.

**Supplementary Figure. 5.**
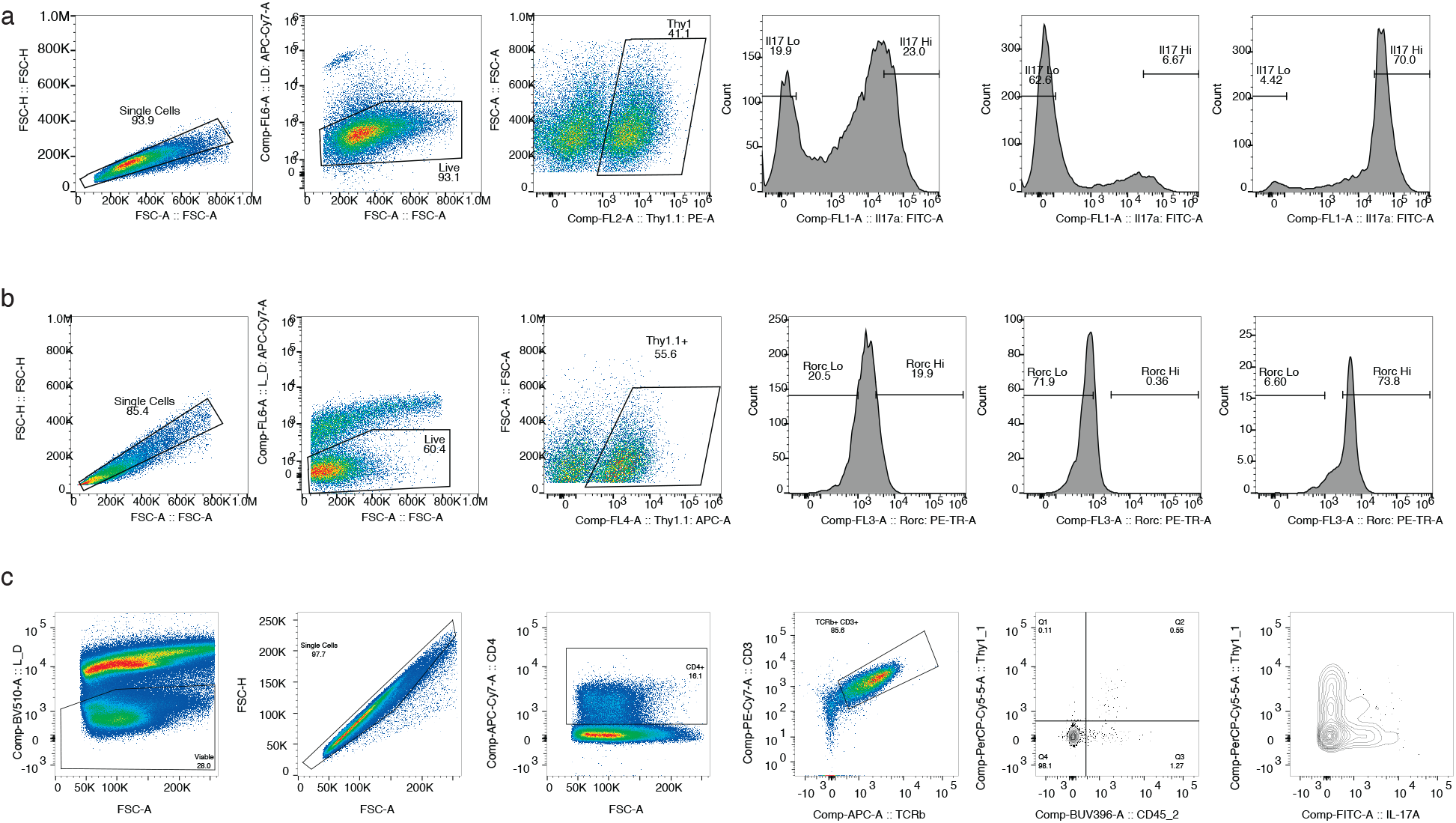
Representative gating strategy for flow cytometry in this study. **a)** Example of Il17a CRISPRi sort for Lymphocytes/Live/Thy1.1+/Il17a-eGFP hi/lo and post-sort checks for Il17a-lo and Il17a-hi bins. The same gating was used for Il17a/f HCR-FlowFish sorting. **b)** Gating strategy for RORγt CRISPRa/i experiments depicting Lymphocytes/Live/Thy1.1+/RORγt hi/lo and post sort checks for RORγt-hi RORγt-lo bins. The same gating strategy was applied for BATF CRISRPa/i. **c)** Gating strategy for in vivo analysis of 7B8-KRAB Th17 cells recovered from the siLP depicting Live/Singlets/CD4+/CD3+ TCRβ+/ CD45.2+ Thy1+ cells.

**Supplementary Figure. 6.**
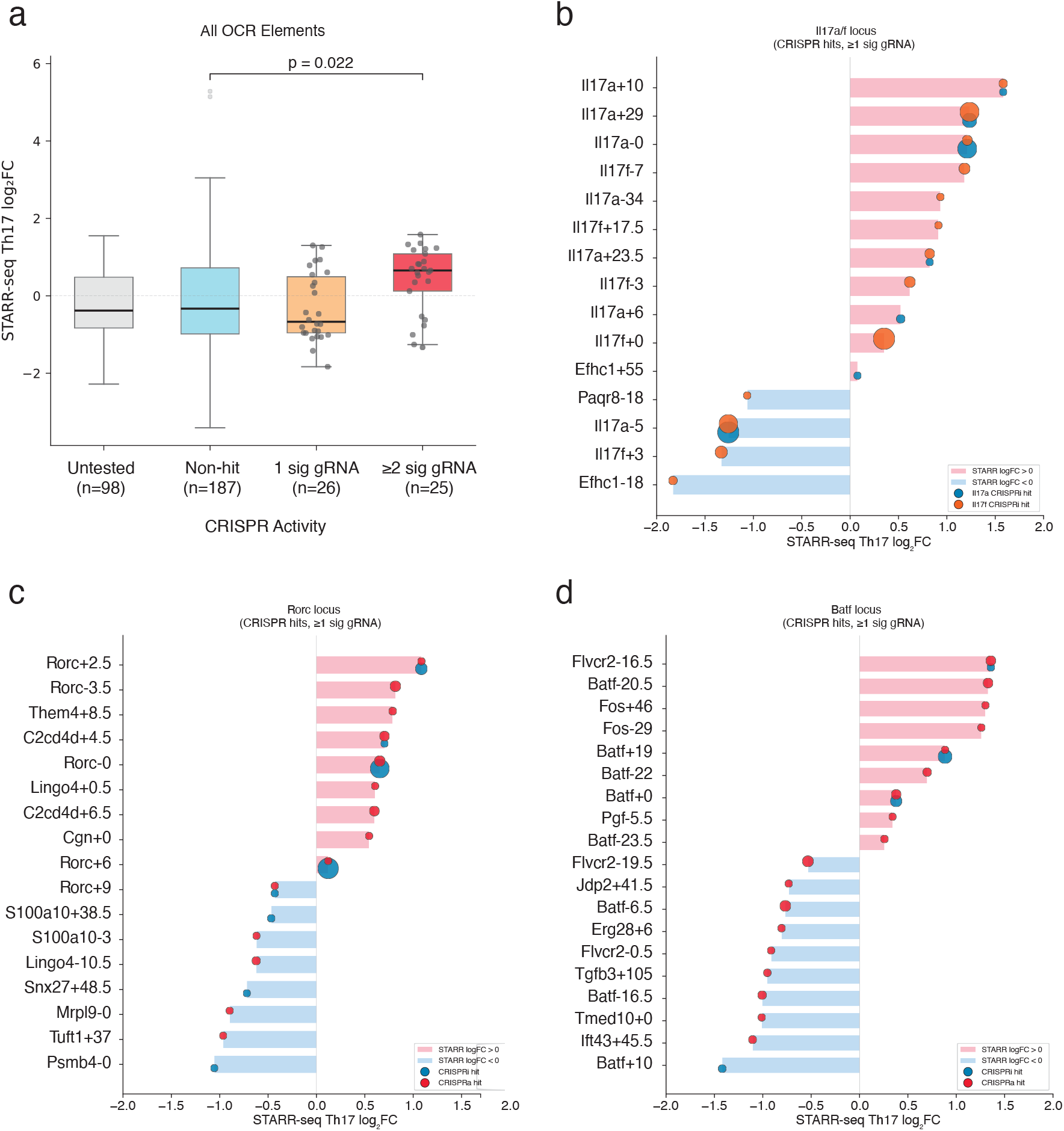
Relationship between STARR-seq and CRISPR functional elements. STARR-seq signal (Log2 FC) at all ATAC-STARR-seq tested OCRs within Batf, Rorc(t), and Il17a/f loci categorized by CRISPR-screen result (untested = no CRISPR gRNA coverage). **b)** Waterfall plot of Th17 cell ATAC-STARR-seq signal (Log2 fold change RNA/DNA) for OCRs with at least 1 significant gRNA in Il17a-CRISPRi (blue circle) or Il17f-CRISPRi (orange circle), **c)** RORγt-CRISPRi (blue circle) and RORγt-CRISPRa (red circle) or **d)** Batf-CRISPRi (blue circle) and Batf-CRISPRa (red circle)

**Supplementary Figure. 7.**
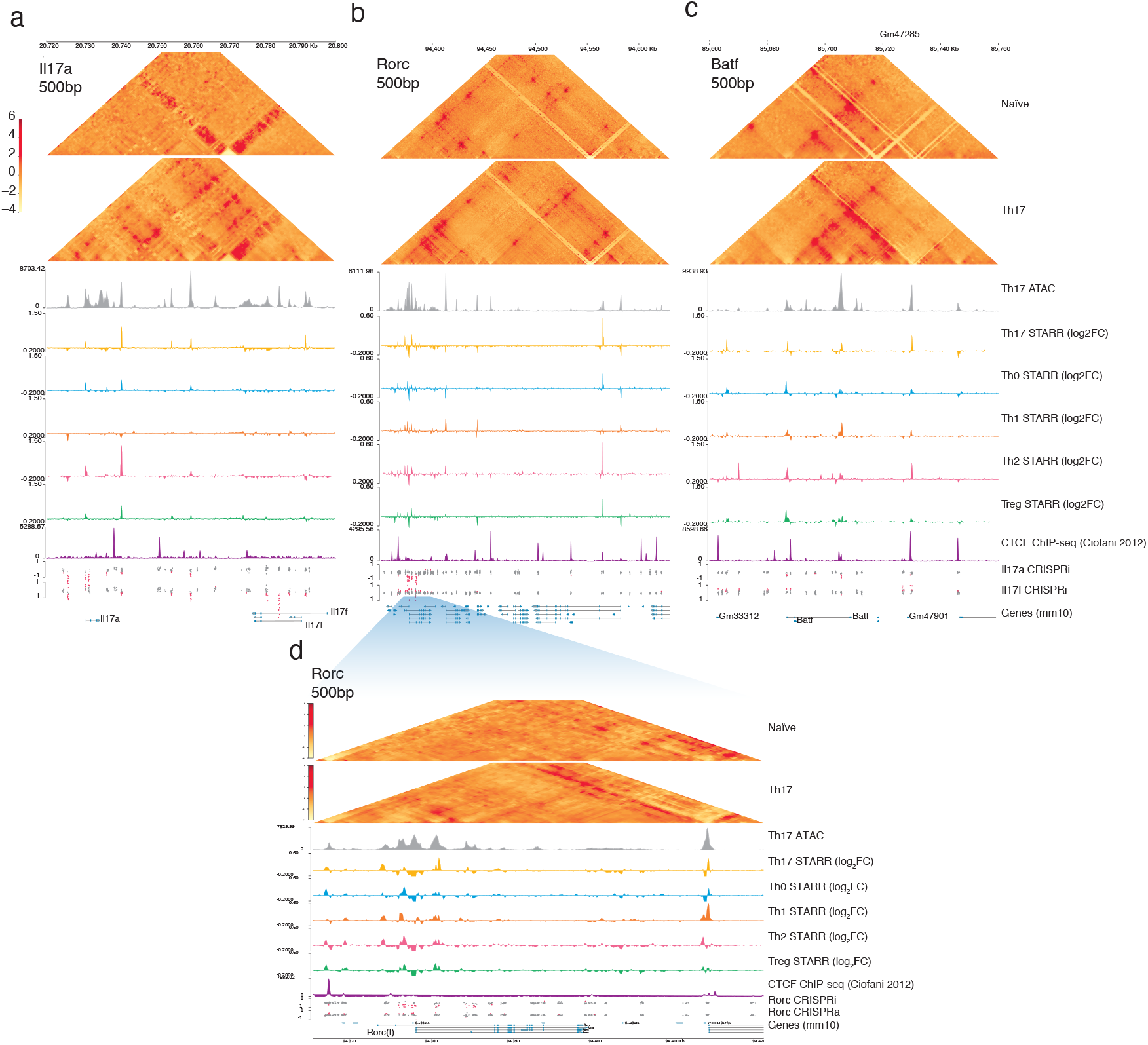
3D Chromatin Architecture of Tnaive and Th17 cells at target loci. Browser track elaborating on chromatin architecture of the **a)** Il17a **b)** Rorc(t) **c)** Batf locus, and **d)** magnified view of the regulatory core region of Rorc(t). Contact heatmaps show RCMC of Tnaive (top) and Th17 (bottom) cells at 500 bp resolution. Track plots include signal from Th17 cell ATAC-seq, Th17, Th0, Th1, Th2 and Treg subset-specific STARR-seq (Log2 fold change RNA/DNA), CTCF ChIP-seq (CPM; Ciofani 2012) and CRISPRa/i screening (Log2 FC hi/lo).

**Supplementary Figure. 8.**
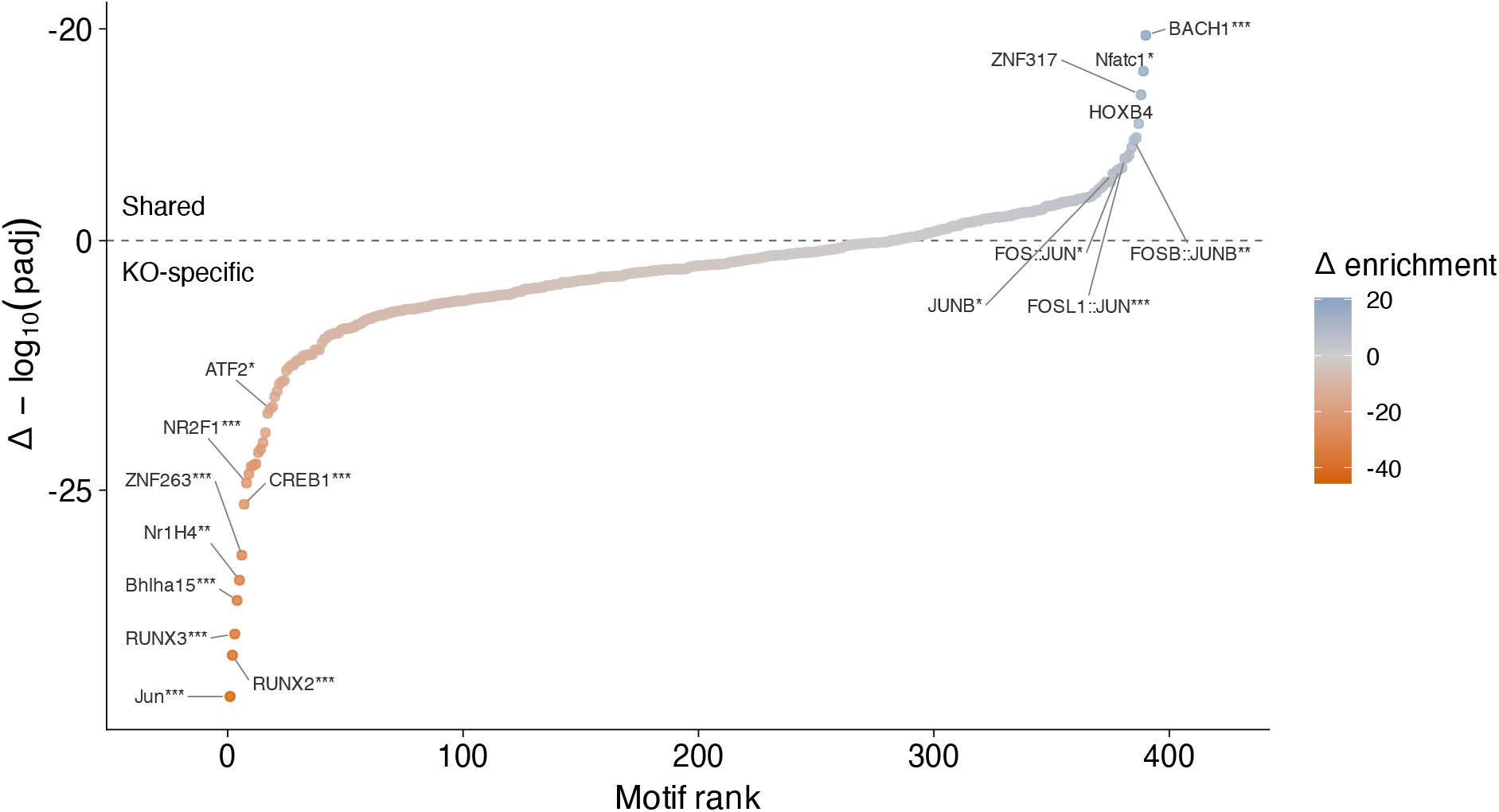
Motif enrichment in condition-specific versus shared differentially accessible OCRs. Differential motif enrichment comparing ATAC-seq peaks with significantly altered accessibility exclusively in Batf-/- relative to control (KO-specific) versus peaks affected in both Batf-/- and Batf +19kb CRISPRi-perturbed (Shared) in vitro polarized Th17 cells (Fisher’s exact test; * FDR < 0.05, ** FDR < 0.01, *** FDR < 0.001).

**Supplementary Figure. 9.**
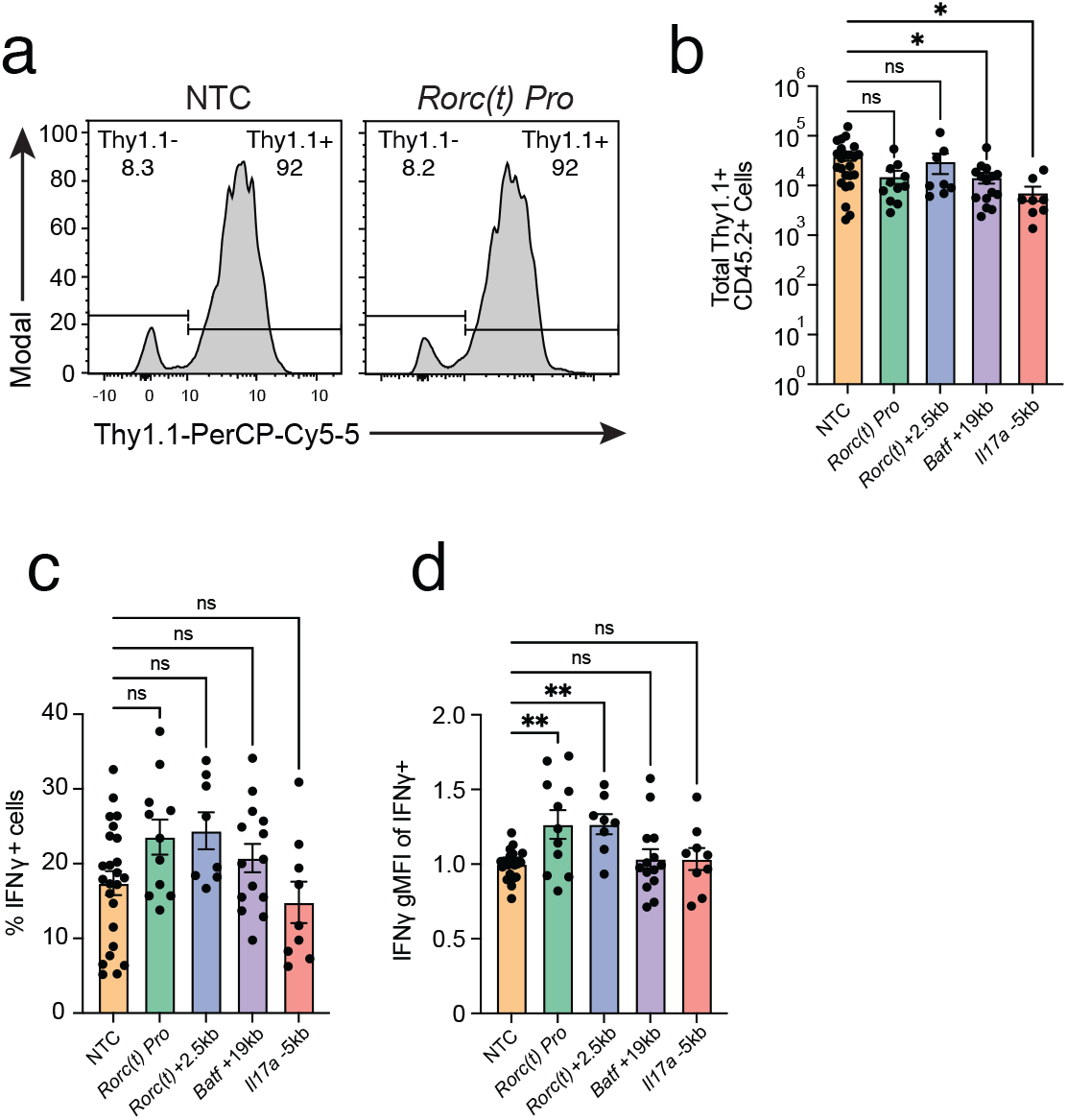
a) 7B8+ Cd4-cre R26LSL-dCas9-KRAB; CD45.2+ CD4+ T cell recovery from siLP is comparable across conditions. **b)** Summary plot of Thy.1.1+ donor cell recovery. **c)** Summary plot of %IFNγ+ 7B8+ Cd4-cre R26LSL-dCas9-KRAB; Thy1.1+ CD45.2+ CD4+ T cells from siLP across conditions. **d)** Summary plot of IFNγ gMFI signal from siLP cells across conditions (** = p<0.01; one-way ANOVA, Dunett’s Test)

## Extended Figure Legends

### Supplementary Tables

**Table 1 Master table of OCRs tested in ATAC-STARR-seq and corresponding Genomic Assays.** Rows indicate individual OCRs (n = 135,897); Columns include genomic coordinates; MACS2 peak calls per cell type (FDR < 0.001); chromatin accessibility classification (shared, unique, combination, or remaining); mean ATAC-seq read depth per cell subset (CPM); ATAC-STARR-seq RNA output read depth per cell type (CPM); and differential enhancer activity statistics from CSAW including raw p-value, Benjamini-Hochberg adjusted FDR, and representative log2 fold change from best test window in CSAW analysis.

**Table 2 Genomic Coordinates of radial plot OCR categories.** Bed coordinates of the top most subset-preferred (r > 0.85) and subset-shared (r < .15) enhancer calls from radial plot in Fig. 1h

**Table 3 Summary of OCRs targeted by CRISPR.** Rows indicating each tested OCR and gRNA per library used in this study

**Table 4 Master Table of CRISPR gRNA**. Rows indicate all gRNA used in this study including genomic location, protospacer sequence, GuideScan output results for library design, Th17 or Th0 cell chromatin-accessibility score, gRNA rank-in-peak, and DESeq2 results by library and CRISPR modality

**Table 5 DESeq2 results table for RNA-seq**. Comparisons for in vitro derived Th17 cells from *Batf* -/- or CRISPRi + *Batf* +19kb gRNA versus non-targeting control (NTC) gRNA.

**Table 6 DESeq2 results table for ATAC-seq OCRs.** Comparisons for in vitro derived Th17 cells from *Batf* -/- or CRISPRi + *Bat f*+19kb gRNA versus non-targeting control (NTC) gRNA.

## Methods

### Mice

All experiments in this study were performed with animal protocols approved by Duke University Institutional Animal Care and Use Committee. All mice were housed in specific pathogen free conditions at Duke University and used between 7-16 weeks of age. For genomics assays (ATAC-seq, RNA-seq, RCMC) and STARR-seq experiments, lineage-defining transgenic reporter mice were used: IFNγ-YFP (Jax stock No:017581), IL-4 GFP (4get-GFP), Foxp3-GFP (Jax stock No:006772), Il17a-GFP (Jax stock No:018472). 4get-GFP mice were generously provided by Dr Irah King (McGill University, Montreal, Canada). For all CRISPR studies, Il17a-eGFP-dCas9- KRAB-CD4-CRE and Il17a-eGFP-dCas9-p300-CD4-CRE were bread in house by crossing dCas9-KRAB (Jax Stock No: 033066) or dCas9-p300 (Jax Stock No: 033065) to Il17a-eGFP and CD4-CRE (Jax stock No:17336). For *in vivo* sgRNA validation studies, the following additional commercially available mouse strains were bred in our facility: 7B8 TCR transgenic (stock 027230, The Jackson Laboratories), CD45.1 (Ly5.1) (stock 002014, The Jackson Laboratories), and *Rag2*^-/-^ *Il2rg*^-/-^ (stock 4111, Taconic Biosciences). All strains were maintained on a C57BL/6J background.

### Isolation of mouse naive CD4^+^ T cells

Spleens and lymph nodes were processed with ammonium chloride potassium (ACK) lysis buffer and filtered with a 0.45µM mesh. For all genomics assays (ATAC-seq, RNA-seq, STARR-seq) and in vivo experiments, CD4 T cells were enriched with Magnisort mouse CD4 T cell enrichment kit (Thermo Cat#8804-6821-74) followed by FACS purification of Tn cells (CD4+CD25- CD62L^hi^CD44^lo^) using either a Beckman Culture Astrios or SONY SH800Z. Fluorescent antibodies for Tn sort panels included CD4-FITC, CD25-eFluor450, CD62L-APC, CD44-PE, fixable viability dye e780; or CD4-PECy7, CD25-PE, CD44-FITC, CD62L-PEcy5, Live/Dead Red. For CRISPRa/i screens, Tn cells were MACS purified to >80% using Magnisort mouse Naïve CD4+ Tcell enrichment kit (Thermo Cat#8804-6824-74).

### Cell Culture and *in vitro* differentiation

All T cell polarizations were done using Genesee tissue-culture plastics pre-coated with anti-Hamster IgG (20 ng/µL in PBS, MP Biomedical Cat#856984) for at least 1 hour at 37°C prior to cell culture.

For all genomic and STARR-seq studies, purified naive CD4+ T cells (Tn) were activated and polarized with subset-specific cytokines simultaneously. Tn cells were seeded at 250k cells/mL. TCR activation was done using hamster-anti-CD3e (0.25 µg/mL), hamster-anti-CD28 (1 µ<u>g</u>/mL). All cultures contained additional blocking antibodies anti-IL4 (2µg/mL, eBiosciences Cat#16- 7041-85) and anti-IFNγ (2µg/mL, eBiosciences Cat#16-7311-85), unless counterproductive (Th1 contained no anti-IFNγ, Th2 contained no anti-IL4). Subset polarization was achieved using subset specific cytokines: Th1 (10 ng/mL IL-12, 10 ng/mL IL-2), Th2 (2 ng/mL IL-4), Treg (5 ng/mL hTGF-β1, 10 ng/mL IL-2), Th17 (10 ng/mL IL-6, 0.3 ng/mL hTGF-β1), and Th0 (10 ng/mL IL-2).

All cultures were performed for 72 hours, except for Th1 and Th2 cells, where cultures were removed from TCR activation at 72 hours and left to rest until 96hr for improved subset conversion and cytokine production.

For CRISPR studies (in vitro and in vivo) naïve CD4 T cells were activated in Th0 conditions for 20-24 hours prior to transduction, then polarized with Th17 specific polarization conditions the next day.

### ATAC-seq

Test fragments were generated using Omni-ATAC-seq as previously described^53^. Briefly, naive T cells were FACS purified from two (Th0, Th1, Th2, Treg) or four (Th17) biological replicates of lineage-defining reporter mice. Naive cells were cultured *in vitro* and polarized to their respective subset as described above. At 72 or 96 hrs, cells from each subset were sort purified into cold PBS with 0.05% BSA using their reporter: Th0 (Foxp3-eGFP⁻), Th1 (IFNγ-YFP⁺), Th2 (IL4-eGFP⁺), Treg (Foxp3-eGFP⁺), and Th17 (Il17a-eGFP⁺). 50,000 cells from each bio replicate (Th0, Th1, Th2, Treg n=2; Th17 n=4) were split into 4 technical replicates for OMNI-ATAC. Cell nuclei were extracted with ATAC-seq resuspension buffer (RSB; 10 mM Tris-HCl pH 7.4, 10 mM NaCl, and 3 mM MgCl2 in water) plus 0.1% NP40, 0.1% Tween-20, and 0.01% digitonin on ice for three minutes. After wash with RSB plus 0.1% Tween-20, chromatin was transposed with 50 µL of a transposition mix containing 25 µL 2× TD buffer, 2.5 μl transposase (Illumina, Cat#20034197), 16.5 μl PBS, 0.5 μl 1% digitonin, 0.5 μl 10% Tween-20, and 5 μl water for 30 minutes with shaking at 37°C. Transposed DNA fragments were purified with Qiagen MinElute kit into 10 µL. 2 µL (20%) from each technical replicate was taken for ATAC-seq library prep. The remaining 8 µL of fragments were pooled within replicates and processed separately for STARR-seq input library generation.

### RNA-seq

Naive CD4 T cells were sort purified and polarized in vitro using our standard culture system. Each subset was polarized for 0 hr, 2 hr, 4 hr, 16 hr, 24 hr, 48 hr, 72 hr. For Th1 and Th2 cells, cultures were expanded to 96hr as described above. For each time point cells were washed twice with cold PBS before being collected as bulk culture. At the final timepoint, an additional sample was collected for each culture following sort purification of the lineage defining reporter gene (Lymphocytes/SingleCells/Live/CD4+/Subset Defining Reporter+ (Th17 = Il17a-eGFP+; Treg = Foxp3-eGFP+; IFNγ = IFNγ-YFP+; Th2 = IL-4 eGFP+; Th0 = Foxp3-eGFP-). Cell pellets were lysed with Qiagen RLT and 1% ß-mercaptoethanol (ß-ME) then processed using a Qiagen RNeasy RNA Extraction Kit. Each library was constructed using Illumina TruSeq Stranded mRNA Library Prep Kit (cat. no. 20020595) following mRNA-capture and sequenced with a Nextseq 2000 P2 kit with 51 bp PE reads. Raw reads were trimmed using Trimmomatic v0.32 - SLIDINGWINDOW:4:20 to remove adapters and low quality read signal. Trimmed reads were aligned to the GRCm38 mouse genome using STAR v2.4.1a, and non-canonical splice junctions were removed (--outFilterIntronMotifs RemoveNoncanonical). Aligned reads were assigned to genes in the GENCODE vM25 comprehensive gene annotation using Subread featureCounts with default settings. Differential expression analysis on feature counts was performed using DESeq2

### ATAC-STARR-seq

For a detailed protocol refer to our protocol on the ENCODE4 portal (Accession #’s: ENCSR576KQE; ENCSR763SPB; ENCSR300TTB; ENCSR945FEC; ENCSR744ZON).

### Insert Library Generation

To maintain high diversity of fragments, the tagmented DNA remaining from OMNI-ATAC were pooled between replicates for each cell type. Pooled fragments were PCR amplified with KAPA HiFi PCR kit (Roche/KAPA, Cat#KK2101) using custom primers containing 5’ homology arms complementary to the 3’UTR cloning region of a modified ORI-STARR-seq vector. The vector was a gift from Alexander Stark (Addgene #99296) and was modified to replace the truncated GFP with cDNA from Thy1.1 (CD90.1).

### Input Library Generation

Inserts were cloned with Gibson assembly into linearized ORI-STARR-Thy1 vector using 10x 20µL 10 nM NEBuilder (New England Biolabs, Cat#E2621X) reactions and a 3:1 DNA insert:linearized backbone ratio. Reactions were incubated at 50C for 2 hours, then pooled and DNA was recovered with overnight EtOH precipitation. Plasmid DNA was resuspended in 10 µL of 10mM Tris-HCl pH 8.0, and split across 4x300 µL aliquots of Endura Duo ElectroMax cells (Lucigen, Cat#60242-1) expanded and prepared in house (See ENCODE protocol). DNA was electroporated using a BioRad GeneMax Pulse Unit in 2mm cuvettes with an exponential decay pulse wave set to 3000 V, 200 Ω, and 25 µF. Cells were recovered with 5mL of recovery media (Lucigen, Cat#80026-1) per cuvette for 1hr at 37C with shaking. All cuvettes were pooled and split across 2x4L Erlenmeyer flasks containing 1L Luria Broth (LB) + 100ng/mL carbenicillin. To maintain high diversity, cells were grown for only 10-12 hours at 37°C with shaking at 220RPM. ATAC-STARR-seq input libraries were recovered from bacteria with MN NucleoBond EF 10000 Giga Prep kit (Macherey-Nagel, Cat#740548) following the manufacturer’s instructions. Plasmids were concentrated to 3 µg/µL in endotoxin free TRIS-EDTA (EF-TE) pH 8.0 and stored at 4°C. To generate the pooled ATAC-STARR library, plasmid libraries from Th0, Th1, Th2, Th17, and Treg STARR-seq input library preps were added together at equal mass.

### T cell Nucleofection of ATAC-STARR-seq libraries

Naïve CD4 T cells were isolated from mouse spleens and lymph nodes by FACS as above. Cells were plated at 4x10^6^ cells per 10mL on anti-hamster IgG (MP Biomedical Cat#856984; 1:50 in PBS) coated 10cm Olympus culture plate (Genesee Scientific cat # 25-202), then polarized with the activating culture conditions described above for each respective subset. At 72 hr (Th0, Th17, Treg) or 96 hrs (Th1, Th2) post-activation, polarized T cells were harvested counted and centrifuged at 80 x g for 10 minutes at 4C before nucleofection. Between 100-200M cells were used per replicate. Approximately 15 × 10⁶ cells per cuvette were resuspended in P3 Primary Cell Nucleofector Solution (Lonza Cat#V4XP-3024) and mixed with ∼36 µg of STARR-seq plasmid library DNA per cuvette (12 µL of 3 µg/µL library). Cells were electroporated using a Lonza 4D- Nucleofector X Unit (program DN-100). Immediately after nucleofection, cells were transferred to pre-equilibrated recovery media. To reduce cell death, complete RPMI was used to recover all subsets, as IMDM has high calcium content, and polarizing cytokines plus blocking antibodies were included without anti-CD3/CD28. Three cuvettes were combined and re plate at high density in an uncoated 10cm plate for 4hr at 37°C. Cells were then replate onto anti-hamster IgG-coated dishes and restimulated with anti-CD3/CD28 restimulation for 2 hr (6 hr total post-nucleofection). Finally, cells were harvest with centrifugation for 10 minutes at 80 x g, and lysed in RLT buffer supplemented with β-mercaptoethanol. Total RNA was isolated using the RNeasy Midi kit (Qiagen, Cat. #75144) and stored at −80°C.

### ATAC-STARR-seq Output Library

STARR-seq RNA output libraries were generated from total RNA isolated from nucleofected T cell subsets. mRNA was purified from approximately 300 µg of total RNA using two sequential captures with four oligo(dT) Dynabead (Invitrogen Cat#61005) columns. Purified mRNA was treated with TURBO DNase (Ambion, Cat#AM1907) for 30 min at 37°C to remove residual plasmid DNA and inactivated with DNase Inactivation Reagent. First-strand cDNA synthesis was performed using SuperScript III reverse transcriptase (Invitrogen, Cat#18080085) with a custom UMI-containing RT primer at 50°C for 2 hr, followed by heat inactivation at 70°C for 15 min and RNase A digested (Ambion Cat#AM2271) for 30 minutes at 37°C. cDNA were purified with 1.4× SPRI beads, and spliced STARR-seq cDNAs were enriched with 8 replicate PCRs per sample using the Kapa Hot-start HiFi 2X master mix (KAPA Cat#: 50-196-5215), the intron junction-spanning PCR (jPCR) forward primer, and the P7 adapter reverse primer (PCR conditions: 15 cycles; 64.5°C annealing). After 1X SPRI bead cleanup, samples were split again to 7 replicate nested PCR reactions with Nextera-indexed i5 forward primer, and P7 adapter reverse primer (PCR conditions: 6 cycles; 64.5°C annealing). Resulting libraries were purified by SPRI bead cleanup (Beckman Coulter, Cat#A63881), quantified by TapeStation, and pooled for sequencing on an Illumina NextSeq 2000 with 50bp PE reads.

### ASTARR-seq differential enhancer activity analysis

Enhancer activity from ASTARR-seq libraries was quantified using a sliding window approach implemented in CSAW^54^ and edgeR^55^. Due to an overabundance of short fragments (<140bpm) in output libraries compared to input, BAM files were filtered to retain only paired-end fragments between 150 and 800 bp in length (Supplementary Fig. 2a), with a minimum mapping quality of 20 and PCR duplicates removed. Reads were counted into 50 bp sliding windows with a 25 bp step size. Windows were restricted to those overlapping the union peak set derived from MACS2^56^ ATAC-seq and input DNA peak calls (FDR < 0.001). Composition biases were corrected using CSAW’s normFactors and 2,500 bp bins. For edgeR differential testing, all cell types (Th0, Th1, Th2, Th17, Treg) and input DNA samples were modeled together using quasi-likelyhood. Contrasts compared each T helper subset against the input DNA. Overlapping significant windows were merged with a tolerance of 0 bp and a maximum merged width of 3,000 bp. Within each merged region, window-level p-values were combined using CSAW’s minimalTests with summit-based abundance weighting (upweightSummit), requiring a minimum of 3 significant windows per region. Multiple testing was adjusted using Benjamini–Hochberg and regions were classified as active with FDR ≤ 0.05.

### Sequence Motif Analysis with gapped k-mer SVM

We trained gapped k-mer support vector machines (gkm-SVM) classifiers^22,23^ using the R package gkmSVM with default parameters (word length L = 10, informative positions K = 6, maximimum mismatches d=3). For each T cell subset, we compared activated (FDR < 0.05, Log_2_ FC > 0), repressed (FDR < 0.05, Log_2_ FC < 0), and inactive (FDR > 0.80). Genomic sequences were extracted from mm10 using BSgenome.Mmusculus.UCSC.mm10 package. Inactive sequences were GC matched for each comparison class (active or repressed), and total sequences per class were capped to 5000 per cell type. Two separate classifiers were trained: active vs inactive, and repressed vs inactive. Classification performance was evaluated using 5- fold cross validation where each fold trained on 80% and held-out 20% then pooled to compute a single AUC per cell type. Top scoring k-mers were annotated against JASPAR 2022 vertebrate TF binding profile database using position-wise PWM scoring and retaining only the best matching TF.

### Subset specificity of ASTARR-seq activity

To classify enhancers by their cell-type specificity across CD4+ T helper subsets, we developed a weighted effect-size metric and projected it into a two-dimensional star-coordinate space. ASTARR-seq enhancer activity was quantified across five conditions (Th0, Th1, Th2, Th17, and Treg), with differential activity assessed using CSAW^54^, keeping only OCRs with activity (FDR < 0.01) in at least one subset. Weighted effect sizes were then computed for each OCR and subset by multiplying absolute Log_2_ fold change by the -log10 FDR to capture both the magnitude and statistical confidence of subset specific activity. Per-region weighted scores were normalized across each subset so that the proportional contributions summed to one.

To visualize the multi-dimensional specificity profile of each enhancer in two dimensions, we used a star-coordinate projection. Each of the five cell types was assigned an equally spaced angle on radial plot (i.e θ = 0, 2π/5, 4π/5, … etc.). For each enhancer, the Cartesian coordinates were computed as: *x* = Σ(*s*_i × cos(θ_i)), *y* = Σ(*s*_i × sin(θ_i)), where *s*_i is the subset effect score and θ_i is the angle assigned to cell type *i*. In this projection, enhancers with STARR-seq activity biased toward a single subset fall near the corresponding spoke of the radial plot, while enhancers with similar activity across all subsets cluster near the origin (Supplementary Fig. 4). For visual clarity, each enhancer was coloured by the subset with the highest effect score. To identify highly subset-specific enhancers for motif analysis, we selected regions with an arbitrary radial distance (*r* > 0.85) from the origin to select elements whose angular position fell closest to the assigned spoke for their dominant cell type (up to 2000 were selected). Shared enhancers with broadly equivalent activity across conditions were identified as those with *r* < 0.10 (up to 5000 were selected).

### CRISPR Screening

#### Library Design

To prioritize regulatory elements involved in Th17 differentiation, we rank ordered the differentially accessible chromatin regions (dATAC) nearest to the most differentially expressed genes (DEGs) when comparing ATAC-seq and RNA-seq from Th17 to Th0 at 72 hours culture in vitro. The top 7000 open chromatin regions were selected (Supplementary Table 3). For targeted screens, all Th17 ATAC-seq peaks detected using MACS2 were selected within the loci Il17a-Il17f (chr1:20,198,461-22,383,378), Rorc (chr3:93,261,362-95,108,248), and Batf (chr12:84,681,315- 86,479,642) (Supplementary Table 3).

SpCas9 guide RNA were selected from a query of these regions against the mm10 Cas9 database using GuideScan 1.0^57^ (Expanded screen) and GuideScan 2.0^58^ (Targeted Screens). sgRNA were then filtered to remove GuideScan specificity scores < 0.2, and repeat strings of GGGGG or TTTT. We further ranked gRNA by Th17 chromatin accessibility by counting the number of ATAC-seq fragments overlapping each gRNA coordinate. Guides within each target region were ranked in descending order of accessibility score, quantified as the mean sum of ATAC-seq fragments (normalized to CPM) overlapping each 20 bp gRNA target using deeptools multiBamSummary across four Th17 or two Th0 biological replicates. The maximum ATAC CPM of the two T cell subset means was taken as the final accessibility score for each guide. To minimize sgRNA overlap and maximize coverage from the OCR summit, guides were accepted in rank order only if they overlapped no previously selected guide by more than 10 bp. Either 10 or 15 non-overlapping guides were retained per target region for the large and targeted libraries, respectively. We designed non targeting control (NTC) gRNA to be 5% of the total library. NTC protospacers were GC content matched to targeting gRNA and did not align mouse genome (bowtie2 [x parameters]. For each gRNA library we added 5-10% of the total as NTCs. The final gRNA row-wise data tables are described in Supplementary Table 4.

gRNA oligo pools were ordered from Twist Bioscience. The 10ng linear template was PCR amplified for 10 cycles in 50 µL reactions using KAPA HiFi (KAPA, Cat #50-196-5215) and primers with homology arms compatible with pooled cloning into the MSCV-mU6-v2Tracr-Thy1.1 gRNA delivery vector. Backbones were linearized with Bbsi-HF digestion with a ratio of 2µg plasmid per 1 µL enzyme. Linear backbone was recovered with .65X Ampure Beads and purified in 10mM Tris. For targeted libraries, we used 5x20 µL 10 nM NEBuilder (New England Biolabs, cat #E2621X) reactions at an insert:backbone ratio of 5:1 for cloning of each targeting library. For our expanded library, we used 20x20 µL 10 nM NEBuilder reactions. Libraries were purified with EtOH precipitation.

Library-containing MSCV viral particles were produced using Platinum-E retroviral packaging cell line (293T cells; Cell Biolabs; Cat#RV-101). Four million 4M PLAT-E cells were seed onto 10cm dishes in RPMI + 10%FBS and pen/strep. After 16 hours, 2.5µg of plasmid was transfected using lipofectamine 3000 (Thermo, Cat#L3000015) following manufactures protocol. Transfected media was changed after 6 hours with RPMI + 10% FBS and virus-containing media was harvest with replacement at 24 hours and 48 hours post transfection. Virus was stored at 4C between collections, then pooled together, centrifuged at 3000 x g for 10 minutes at 4C to remove debris. Viral supernatant was then concentrate using RetroX (Takara, Cat#631456) following manufactures instructions, concentrated to 50x aliquots and stored in liquid nitrogen.

#### T cell retroviral transduction

Tn cells were isolated and purified from the spleen and lymph nodes of *Cd4:Cre+/dCas9-p300+* or *Cd4:Cre+/dCas9-KRAB+* mice using MACS as described above. 2.5M naïve T cells were seed on 6 well plate in Th0 activating conditions for 24 hours. Cells were transduced with murine stem cell virus (MSCV) and gRNA libraries or single gRNA for validation and 6.66 ng/mL polybrene using spin transduction for 2 hours at 2400 RPM and 30°C. For CRISPR screening, viral titres were estimated using flow cytometry, and transduction was performed at low multiplicity of infection (MOI; ∼0.3), and transduction rates were consistently between 30-50%. For validation studies MOI was not estimated and transduction rates ranged from 20-60% per gRNA viral prep. 12-16 hours post spinfection, viral supernatant was replaced with Th17 polarizing media and cells were cultured for 72 hours.

#### *In vitro* CRISPR screening with Flow Cytometry

In this study multiple reporters were applied: Il17a locus-reporter transgenic mice (Figure 2), intercellular protein staining (Figure 2, 4, 5), and HCR-FlowFish (Figure 3). For all steps, cells were first enriched via MACS using Thy1.1 (CD90.1) positive selection kit (StemCell Cat#18958). Thy1+ cells were stained with Fixable Viability Dye (eFluor 780; eBioscience Cat#65-0865-14; 1:1000) and anti-Thy1.1 APC (StemCell, Cat#60024AZ, clone OX-7; 1:400) or Thy1.1-PE (StemCell, Cat#60024PE, clone OX-7; 1:400) or Thy1.1-PerCP-cy5.5 (Thermo, Cat#45-0900-82, clone HIS51; 1:300) as extracellular protein stain dependent on flow panel. Cells were then processed for sorting one of three ways:

#### Intracellular Fluorescent Reporter

Live cells were prepared directly for FACS.

#### Intranuclear Transcription Factor Staining

Prepared cells were fixed and permeabilized using the Foxp3/Transcription Factor Staining Buffer Set (eBioscience; Cat#00-5523-00) according to the manufacturer’s instructions. Fixed and permeabilized cells were stained with fluorescent primary antibodies diluted in permeabilization buffer for 30 min at room temperature in the dark per their respective screens. Rorc(t): anti-RORγt PE (eBioscience, Cat#12-6981-82,clone B2D; 1:200), Batf: anti-BATF PE (BD Bioscience, Cat#564503, clone S39-1060; 1:200).

#### HCR-FlowFISH

Single-cell gene expression was quantified using HCR-FlowFISH as previously described^59,60^. Briefly, cells were fixed in 4% formaldehyde in PBST for 1 hr at room temperature, washed, and permeabilized in 70% ethanol at 4°C. Cells were pre-hybridized in probe hybridization buffer containing 30% formamide and 10% dextran sulfate, then incubated overnight at 37°C with Il17a and Il17f HCR probe sets (Molecular Instruments) at a final concentration of 4 nM. Signal was amplified with fluorescently labeled hairpins for three hours at room temperature. All wash steps were performed in low bind tubes at 800 x g with no brake applied to centrifuge deceleration to maximize cell retention.

#### Fluorescence Activated Cell Sorting

Cells were FACS purified into bins of top and bottom signal distribution. For expanded screens, we sort 2.5M cells from two replicate pools of mice. For targeted screens, we took at least 500k cells from three replicate pools of mice. For Il17a and IL17f screens, the reporter distribution was bimodal; therefore, we sort from the top 20% of Il17a-eGFP+ or Il17a+ cells, and the bottom 20% of negative signal (Supplementary Figure 5a). For BATF and RORγt screens, we sort from the top and bottom 20% of fluorescent signal (Supplementary Figure 5b). All sorts were performed using a Sony SH800Z or Beckman Coulture Astrios. The final gating strategy was Lymphocytes/ Single Cells/ Live/ Thy1^+^/ and high / low populations of: Il17a-eGFP^hi/lo^ or *Il17a^hi^*^/lo^ or *Il17f^hi^*^/lo^ or BATF^hi/lo^ or RORγt^hi/lo^)

#### gRNA Recovery and output CRISPR screening library prep

Genomic DNA was recovered from sorted bins using Zymo Quick-DNA mini prep (Zymo Cat # D3025) following manufactures instruction, including all additional washes and elution steps. Final libraries were PCR amplified using NEB q5 2X Master Mix in 100 µL PCR reactions using a maximum of 1 µg of gDNA template and custom primers designed to amplify the gRNA protospacer with Illumina TruSeq adaptors from the MSCV cassette. Libraries were sequenced on an Illumina NextSeq 2000 with 21 bp single end reads.

#### gRNA processing and alignment

Sequencing reads from CRISPR screen libraries were trimmed to the 20 bp protospacer sequence using Trimmomatic (v0.32; CROP:20)^61^. Trimmed reads were aligned to a custom reference genome constructed from the corresponding gRNA library by using Bowtie2 in end-to- end mode and with flags --very-sensitive -a --norc. Aligned reads were converted to sorted, indexed BAM files using samtools, and per-guide read counts were extracted with samtools idxstats. Differential gRNA abundance between high and low fluorescence-sorted cell populations was assessed using DESeq2^62^.

#### Region Capture Micro-C

We sort purified 25M Naïve CD4 T cells (CD4+CD25-CD62L^hi^CD44^lo^) or in vitro derived Th17 Il17a-eGFP+ cells for input into RCMC. We followed the RCMC protocol from Hansen lab^63^ with few modifications. Specifically, we empirically determined MNase (Worthington Biochemical, Cat #LS004797) chromatin fragmentation conditions for Naïve and Th17 cells to be 2U MNase for 12 minutes at 37C per 1M cells. In addition, hybridization temperature for probe-based capture of Micro-C libraries proved to be critical, so heat-blocks and thermocyclers were calibrated and verified using a temperature gauge before proceeding.

Region-capture probes were designed by Twist Bioscience to multiplex-capture the Il17a/f (chr1:20198461-22383378), Rorc (chr3:93261362-95108248), and Batf (chr12:84681315-86479642) loci using end-to-end tiling with 80-120mer non-overlapping probes, and low off-target stringency settings. 80.9% of the 5.8 Mbp total region was covered successfully. A single hybridization was performed for all three libraries, per sample.

#### RCMC Data Processing

Region-capture Micro-C (RCMC) libraries targeting the Il17a/Il17f (chr1:20,198,461-22,383,378), Rorc (chr3:93,261,362-95,108,248), and Batf (chr12:84,681,315-86,479,642) loci were sequenced on an Illumina NextSeq using paired-end (PE) 50bp reads. Reads were aligned to the mm10 mouse reference genome (ENCODE no-alt analysis set) using Bowtie2 --very-sensitive- local --reorder. Aligned reads were parsed into contact-pairs using pairtools parse --min-mapq 2 --walks-policy mask. Pairs were sorted with pairtools sort and deduplicated with pairtools dedup --max-mismatch 1 --mark-dups. Deduplicated pairs were split into BAM and pairs formats using pairtools split, then filtered with pairtools select to retain only read pairs in which both mates mapped within one of the three target loci. Filtered pairs were sorted and indexed using pairix.

Contact matrices were first generated at 50 bp resolution using cooler cload pairix with genome- wide bins defined by the mm10 chromosome sizes. To maximize sequencing depth for visualization of baseline chromatin architecture at targeted loci, RCMC data from wild-type (untransduced) and NTC-transduced Th17 cells were pooled. Biological replicate Tnaive RCMC contact matrices were similarly pooled. Pooled matrices were used for contact map visualization in the main and Supplementary figures. Quantitative comparisons of contact frequency between NTC and CRISPRi-perturbed conditions used only the NTC replicate data. To visualize each target locus, bins and pixels were extracted from the genome-wide 50 bp cooler using cooler dump, re-indexed to produce locus specific coordinates, and loaded into locus-specific cooler files with cooler load. Multi-resolution contact matrices (.mcool) were then generated with cooler zoomify at resolutions of 50, 100, 250, 500, 1,000, 2,000, 5,000, and 10,000 bp, with iterative correction (ICE) balancing applied at each resolution (--balance --min-count 1). Multi-resolution contact matrices were visualized in Python using the coolbox API and matplotlib. ICE balanced, observed/expected (O/E) normalized matrices were displayed at 250 to 500 bp resolution.

#### Aggregate loop signal quantification

Aggregate contact signals were computed for three specific interactions in the Batf locus: promoter to Batf +19kb (P-E1), promoter to Flvcr2 -16.5kb (P-E2), and Batf +19kb to Flvcr2 - 16.5kb (E1-E2). Enhancers were located at chr12:85,705,664 and chr12:85,729,327. For each interaction, the mean O/E value was calculated within a ±2-bin (±1 kb) window centered on the corresponding target for both NTC and gRNA transduced samples.

#### In vivo Th17 cell differentiation model

Naïve CD4^+^ T cells were FACS-sorted from spleen and lymph nodes of adult 7B8 TCR transgenic *Rosa26*^dCas9-KRAB/WT^ *Cd4*-Cre^+/−^ CD45.2 mice and cultured under Th0 conditions (without IL-2) in complete IMDM on anti-hamster-coated plates (0.8 × 10^6^ cells per well). Cells were transduced 20-24h post activation with MSCV retrovirus containing sgRNA constructs (non-targeting control or gene-targeting sgRNA). Spin transduction (spinfection) was performed using viral supernatant supplemented with polybrene (6.66 µg ml^-1^) and centrifugation in a parafilm-sealed plate at 2,400 rpm for 2hrs and 30 °C. Activating Th0 media was added back and cells were cultured for an additional 24 h. Thy1.1^+^ cells were then FACS-sorted and 2-5 × 10^5^ cells were adoptively transferred by retro-orbital injection into sex-matched SFB-free CD45.1 (Ly5.1) recipient mice. The following day, recipient mice were orally gavaged with 50mg of fecal slurry from SFB-colonized *Rag2*^-/-^ *Il2rg*^-/-^ mice to induce intestinal Th17 cell differentiation. Eight to ten days after adoptive transfer, small intestinal lamina propria lymphocytes (siLP) were isolated as previously described^64^, and analyzed by flow cytometry following *ex vivo* stimulation and intracellular staining for IL-17A, IFN-γ, Foxp3, and RORγt. Donor-derived Th cells were identified as CD45.2^+^ Thy1.1^+^ CD4^+^ CD3^+^ Foxp3^-^ TCRβ^+^ lymphocytes.

## Data Availability

All raw and processed genomics datasets are available on GEO: GSE342109 and on the ENCODE portal: https://www.encodeproject.org/carts/siklenka2026/. CTCF ChIP-seq data is available from Ciofani et al., 2012^3^

## Code Availability

All code used for analysis is available on GitHub: (https://github.com/ksiklenka/Siklenka2026/) . Additional code available upon request.

## Acknowledgements

We thank the Duke Cancer Institute flow cytometry core, Duke Large Animal Research staff, and Duke sequencing core for supporting the experiments related to this work. Dr Irah King (McGill University, Canada) for the Th2 reporter mouse. Dr Alexander Stark for the original ori-STARR-seq backbone. Dr Greg Wray for careful reading and suggestions during writing of the manuscript.

This work was funded by NIH UM1 1UM1HG009428-01 to T.E.R., M.C., C.G, and G.C.; and R01 AI194223 to M.C. M.E.P. was supported by F31 AI152457 and N.U.M. was supported by F31 AI181082.

## Author contributions

KS, TER, CG, GC, MC conceptualized the project; KS, CZ, LL, performed the functional genomics screens and in vitro validation assays; MEP, NUM performed the in vivo validation experiments; KS performed the primary data analysis and integration; AB, RV supported bioinformatics analysis including CRISPR library design. KS wrote the manuscript with input from all authors. TER, CG, GC, MC funded the study, TER, MC supervised the study

